# Buffering of genetic dominance by allele-specific protein complex assembly

**DOI:** 10.1101/2022.11.17.516863

**Authors:** Mihaly Badonyi, Joseph A Marsh

## Abstract

Protein complex assembly often begins while at least one of the subunits is still in the process of being translated. When such cotranslational assembly occurs for homomeric complexes, made up of multiple copies of the same subunit, this will result in complexes whose subunits were translated off of the same mRNA in an allele-specific manner. It has therefore been hypothesised that cotranslational assembly may be able to counter the assembly-mediated dominant-negative effect, whereby the co-assembly of mutant and wild-type subunits “poison” the activity of a protein complex. Here, we address this, showing first that subunits that undergo cotranslational assembly are much less likely to be associated with autosomal dominant relative to recessive disorders. Moreover, we observe that subunits with dominant-negative disease mutations are significantly depleted in cotranslational assembly compared to those associated with loss-of-function mutations. Consistent with this, we also find that complexes with known dominant-negative effects tend to expose their interfaces late during translation, lessening the likelihood of cotranslational assembly. Finally, by combining protein complex properties with other protein-level features, we trained a computational model for predicting proteins likely to be associated with dominant-negative or gain-of-function molecular mechanisms, which we believe will be of considerable utility for protein variant interpretation.

## Introduction

Almost half of the proteins with experimentally determined structures interact with other copies of themselves to form homomeric complexes (Bergendahl and Marsh 2017), and more than one-third of heteromeric complexes with known structures contain sequence-identical repeated subunits (Marsh et al. 2015). Considering the human proteome, about one-fifth of proteins have been detected to cotranslationally assemble in a simultaneous fashion (Bertolini et al. 2021), whereby two subunits interact while still being translated on the ribosome. Cotranslationally assembling homomers are thought to predominantly undergo cis-assembly, yielding allele-specific complexes (Bertolini et al. 2021; Gilmore et al. 1996; Mrazek et al. 2014; Redick and Schwarzbauer 1995). Repeated subunits in a heteromeric complex are also more likely to have come from the same transcript, especially when their assembly is seeded by cotranslational assembly, which reduces the chance of a single complex containing subunits from two alleles. We are beginning to understand which structural properties are important for or predispose subunits to undergo this mode of assembly. Thus far, these include N-terminally exposed interface residues (Bertolini et al. 2021; Kamenova et al. 2019; Natan et al. 2018; Shiber et al. 2018), a large interface area (Badonyi and Marsh 2022), a high alpha helix content and presence of coiled coil motifs (Bertolini et al. 2021) or domain invasion motifs (Seidel et al. 2022). Although many studies have provided insight into the functional and evolutionary aspects of cotranslational assembly, our understanding of the genetic impact of its allele-specific nature is lacking.

Previously, a potential genetic consequence was proposed on theoretical grounds (Natan et al. 2017; Nicholls et al. 2002; Perica et al. 2012). According to the hypothesis, cotranslational assembly should reduce the likelihood that a dominant-negative (DN) mechanism of disease pathogenesis will be observed. A DN effect occurs when expression of a mutant allele disrupts the activity of the wild-type allele (Herskowitz, 1987; Veitia et al., 2018), causing disproportionate function loss and thus a dominant mode of inheritance. Observational evidence has long suggested that DN effects are common in homomers (Veitia, 2007), most probably because incorporation of a mutant subunit into a complex along with wild-type subunits is enough to “poison” the complex. This assembly-mediated DN effect can lead to a reduction in functional activity exceeding the 50% that would be expected for a simple heterozygous loss-of-function (LOF) mutation. However, cotranslational assembly can give rise to complexes whose subunits are allele specific, i.e. homogenous wild-type or homogenous mutant, potentially buffering the deleterious effect of an otherwise DN mutation (**Figure 1**). This ability of cotranslational assembly can be referred to as its “buffering capacity” against DN mutations.

**Figure 1.**
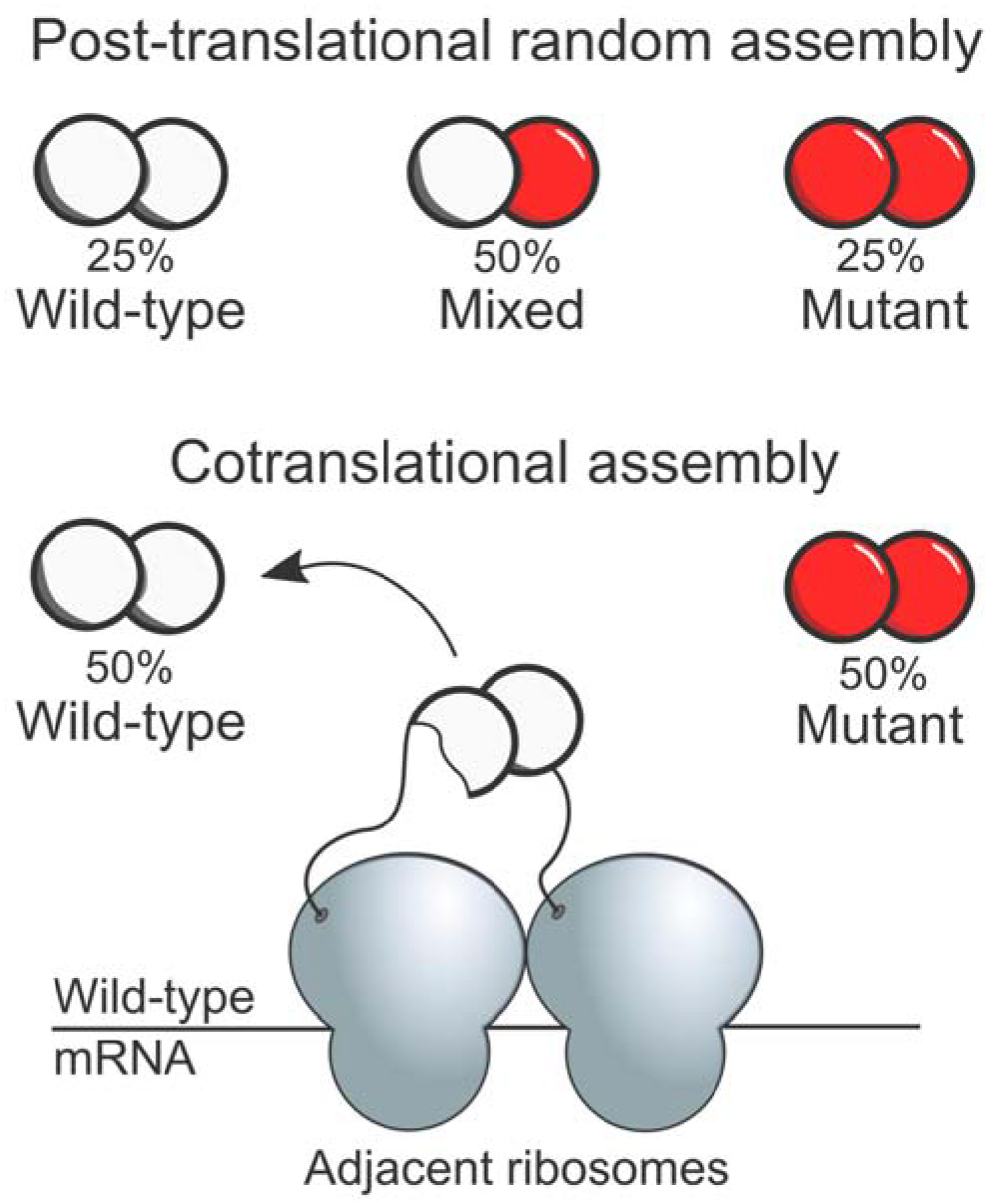
Consequence of allele-specific protein complex assembly Consider a homodimer with one allele of its gene carrying a heterozygous mutation with dominant-negative (DN) properties. When complex assembly occurs after the subunits have been fully translated and folded (post-translational random assembly), the maximum entropy configuration of subunits dictates that mixed complexes will make up half of all complexes. This means that homogenous wild-type and mutant complexes form only 25% of the time. However, when the homodimer cotranslationally assembles, both wild-type and mutant complexes will form independently of one another in an allele-specific manner, increasing the ratio of fully functional complexes to virtually 50%.

Two molecular phenomena may complicate the detectability of such buffering capacity. First, peri-translational assembly (Natan et al., 2017) should make it more likely that post-translationally assembling subunits will also be allele specific, providing that the rate of subunit synthesis outpaces the diffusional force driving the subunit away from the transcript. This effect is likely to be more common in highly abundant proteins, whose transcripts have high ribosome densities and are translated more efficiently than those of lowly abundant proteins (Riba et al., 2019; Schwanhäusser et al., 2011). Second, the phenomenon of subunit exchange (Tusk et al., 2018) may increase the entropy of subunit stoichiometry post-assembly via shuffling of wild-type and mutant subunits, resulting in proportions expected from random post-translational assembly. However, because proneness to subunit exchange is determined by the dissociation constants of the subunits involved, it should be less likely to occur between cotranslationally assembling subunit pairs, which tend to have larger interfaces (Badonyi and Marsh 2022) and thus higher binding affinities.

Heterozygous mutations in genes associated with both recessive and dominant disorders are expected to be enriched in DN mechanisms. First, DN mutations in recessive genes often lead to disease that phenocopy the recessive disorder, as is the case with missense variants in the inositol triphosphate receptor (ITPR1) implicated in Gillespie syndrome (Gerber et al. 2016; McEntagart et al. 2016) or those in the retinoid isomerohydrolase (RPE65) linked to retinitis pigmentosa (Bowne et al. 2011; Thompson et al. 2002). Second, recessivity is an indication that the product of the gene is haplosufficient, i.e. half the gene dose upon a heterozygous null mutation is sufficient to maintain the normal phenotype. Exceptions may be cases where the recessive mutation is hypomorphic, that is, partial LOF, and ultimately results in suboptimal amounts of the protein.

Closely related to the DN effect is a special case of gain-of-function (GOF) mutations. At the protein-level, the phenotypic effect of GOF mutations is the consequence of the mutant protein doing something differently than the wild-type protein, although there is ambiguity in the literature around how this is interpreted. A simple case of GOF is the S810L mutation in the mineralocorticoid receptor (MR), which leads to early-onset hypertension as a result of certain steroid antagonists becoming potent agonists (Geller et al. 2000). However, formation of mixed wild-type:mutant complexes can lead to GOF in a similar manner to the DN effect, but instead of the mutant blocking the activity of the wild-type, the GOF is conferred to the whole complex. This is the case with the L171R mutation in the G protein-activated inward rectifier potassium channel 2 (KCNJ6) implicated in the Keppen-Lubinsky syndrome, which acts by reducing potassium selectivity allowing sodium and calcium ions to pass the channel (Horvath et al. 2018). The precise terminology for this mechanism is the assembly-mediated dominant-positive effect (Backwell and Marsh 2022), which, similar to the assembly-mediated DN effect, may also be subject to genetic buffering by allele-specific assembly.

All things considered, cotranslational assembly may counter the DN effect, which can have an appreciable impact on how genes with inherited and *de novo* missense variants are prioritised in clinical sequencing pipelines. To examine the question, we used a set of cotranslationally assembling proteins determined by disome-selective ribosome profiling (Bertolini et al., 2021) and formulated two hypotheses based upon the above lines of thought: 1) Using dominant disease inheritance as a proxy for the DN effect, homomers and repeated subunits with an autosomal dominant disease inheritance pattern should be depleted in cotranslationally assembly relative to autosomal recessive genes; and that 2) within genes with autosomal dominant inheritance, protein subunits with known DN disease mutations should have the lowest rate of cotranslational assembly. Here, we show that both hypotheses are upheld. Moreover, examination of the structural properties of complexes associated with DN mutations shows that their interfaces are exposed relatively late during translation, which should strongly disfavour cotranslational assembly. Finally, using a knowledge-based approach, we train a regression model to prioritise genes whose mutations are expected to be associated with non-LOF mechanisms. We hope that our work will be of interest to clinical geneticists and accelerate the prediction and discovery of variant-level molecular mechanisms.

## Results

### Autosomal dominant genes are depleted in cotranslationally assembling subunits

We start with a set of 9,053 human proteins, of which 6,562 (72%) physically interact with copies of themselves to form homomeric complexes. The remaining 2,491 (28%) are repeated subunits, meaning that they are present in heteromeric complexes in more than one copy. Both types of proteins have the potential to be associated with assembly-mediated DN or dominant-positive effects, as the mutant and wild-type proteins can co-assemble within the same complex. We obtained genetic inheritance modes from the OMIM database (Amberger et al. 2015) and defined a gene as autosomal dominant (AD) if it had disease mutations inherited in an AD pattern or as autosomal recessive (AR) if it had mutations inherited exclusively in an AR pattern.

An example of a homomer that causes autosomal dominant disease is the ferritin light chain (FTL) complex, which stores and transports iron in a readily available form. Numerous mutations in the *FTL* gene have been linked to neurodegenerative disorders associated with iron accumulation in the brain (Curtis et al. 2001). The frameshift mutation F167SfsX26 in particular (**Figure 2A**) replaces a C-terminal short helix with a stretch of disordered residues, which is thought to have a DN effect by creating large pores in the complex that interfere with its ability to store iron as a solid mineral (Jin et al. 2001). An example of a repeated subunit is found in the GABA_A_ receptor (**Figure 2B**), with both the alpha-1 (GABRA1) and beta-2 (GABRB2) subunits being present in two copies in the complex, an important ionotropic receptor of the central nervous system. The mutation T287P is located in the second transmembrane helix of GABRB2 that forms the inner lining of the channel. It has been proposed that the mutation exerts a DN effect by interfering with the trafficking of the wild-type and reducing the chloride ion current upon activation, ultimately contributing to a developmental and epileptic encephalopathy phenotype (Ishii et al. 2017).

**Figure 2.**
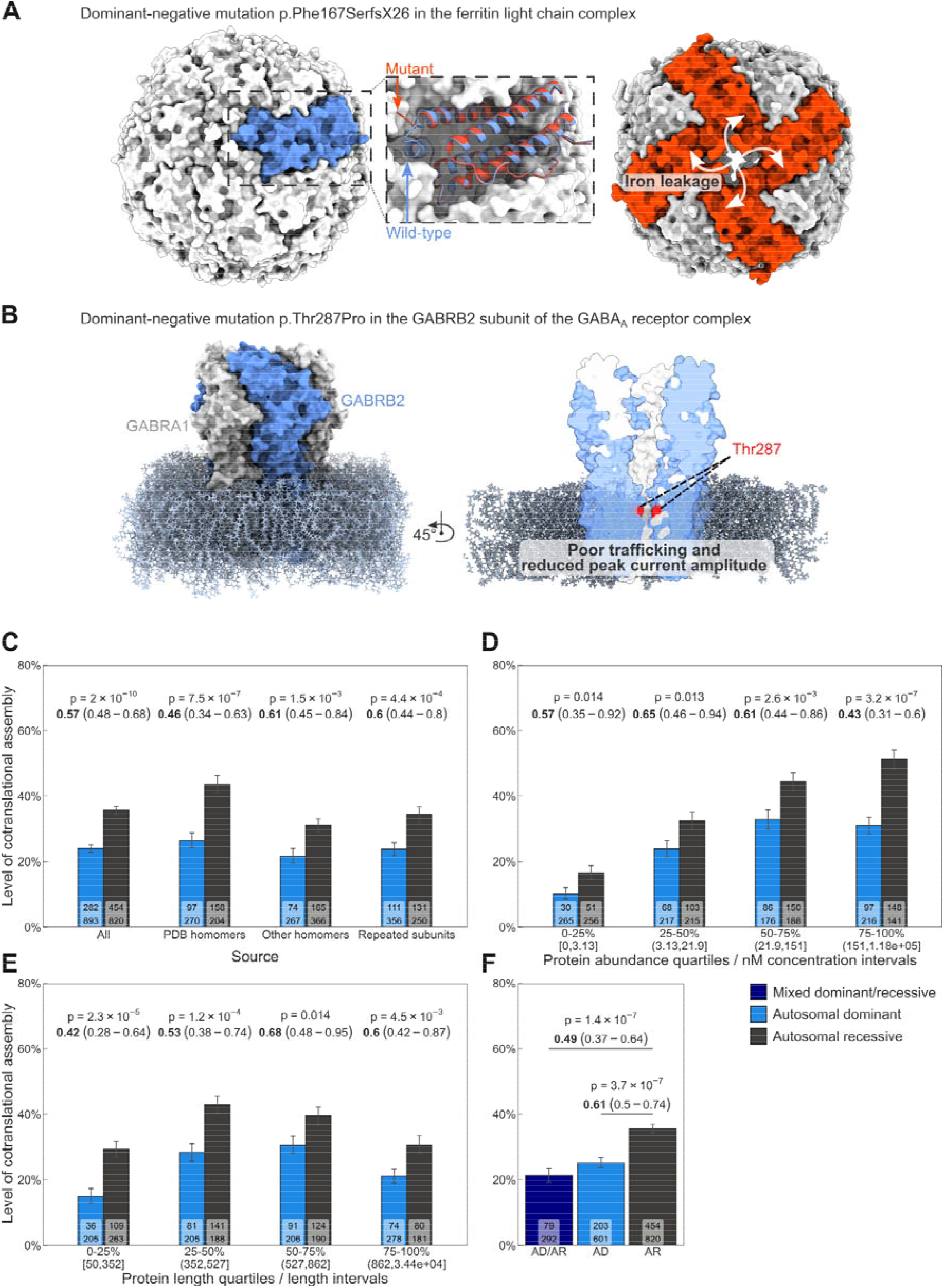
Autosomal dominant genes are depleted in cotranslationally assembling subunits (**A**) Structure of the wild-type ferritin light chain complex (PDB ID: 2ffx) and the F167SX26 mutant (4v6b). (**B**) Structure of the GABAA receptor (6×3s) and location of the residue T287. (**C**) The level of cotranslationally assembling homomers and repeated subunits among autosomal dominant vs autosomal recessive genes subset by subunit source (see **Methods**). The hypergeometric p-value and the odds ratio (OR, in bold) and its confidence interval of the respective comparison is displayed above the bars. In the label on the foot of each bar, the number on top represents the count of cotranslationally assembling subunits and below that is the count of all other subunits. (**D**) The level of cotranslational assembly split into protein abundance quartiles. Each bin corresponds to 25% of proteins by count and the concentration intervals measured in nM are displayed in brackets. (**E**) Same as (**D**) but the bins represent protein length quartiles. Intervals correspond to length in amino acids. (**F**) The level of cotranslationally assembling homomers and repeated subunits among genes with mixed autosomal dominant and recessive inheritance and those with exclusively either one or the other. A Holm-Bonferroni correction was performed on the p-values sampled from the hypergeometric distribution.

Our first hypothesis was that AD genes encoding proteins that assemble into homomers or repeated subunits of heteromers should be depleted in cotranslational assembly relative to AR genes, because subunits with DN mutations should be less likely to undergo cotranslational assembly. While we acknowledge that using AD inheritance as a proxy for the DN effect is a strong simplification, we expect that assembly-mediated DN and dominant-positive effects account for a substantial fraction of AD disorders in these proteins (Bergendahl et al. 2019). We found that 24% of AD subunits cotranslationally assemble compared to 35.6% with an AR inheritance (p = 2 × 10^−10^; hypergeometric test). As a measure of the strength of the relative difference between AD and AR groups, we calculated the odds ratio (OR). To put OR into context using the overall trend, OR = 0.57 can be interpreted as a randomly selected AD homomer or repeated subunit being almost half as likely to cotranslationally assemble as a randomly selected AR subunit. In **Figure 2C**, we show this analysis grouped by the three main sources of the subunit data, which are homomers with experimentally characterised structures (PDB homomers), homomers with non-structural evidence (other, which includes SWISS-MODEL homology models and evidence for homo-oligomerisation from different databases), and repeated subunits of heteromers (see **Methods**). The results show that PDB homomers exhibit the strongest effect (OR = 0.46, p = 7.5 × 10^−7^) followed by repeated subunits (OR = 0.6, p = 4.4 × 10^−4^) and other homomers (OR = 0.61, p = 1.5 × 10^−3^). We speculate that the stronger trend in PDB homomers is due to their enrichment in biologically important homomeric interfaces, which makes a higher fraction of this group amenable to the buffering of assembly-mediated effects. Nonetheless, these results demonstrate that, despite a small variation in effect size, the trend is homogenous across the groups.

We considered a range of confounding variables. Both cotranslational assembly and its detection may be biased by abundance because highly abundant proteins tend to have a higher ribosome density (Riba et al., 2019), which facilitates cotranslational assembly as well as yields more ribosome footprints that can be subjected to sequencing. Notably, AD and AR homomers and repeated subunits have similar median abundance (**Figure S1A**, p = 0.658; Wilcoxon rank-sum test). We binned subunits based upon their approximate intracellular concentration into quartiles, ranging from ∼0.005 nanomolar to ∼180 micromolar, and visualised the differences in the level of cotranslational assembly between AD and AR subunits (**Figure 2D**). Whilst the trend appears strongest in the bin of highest abundance (top 25%, OR = 0.43, p = 3.2 × 10^−7^), it is significant across all quartiles. Because the detection of cotranslationally assembling proteins was performed in HEK293 cells (Bertolini et al.

2021), we repeated the analysis using active ribosome-protected fragment counts specific to HEK293 cells (Clamer et al. 2018), which mirrored the results from the abundance analysis (**Figure S1B**).

Protein length should also be controlled for because AD homomers and repeated subunits are significantly longer than AR (**Figure S1C**, p = 7.1 × 10^−10^; Wilcoxon rank-sum test). Using the same approach, we split the subunits by their length into four quartiles and found that all four bins follow the trend to a statistically significant extent (**Figure 2E**).

Homomers assemble into symmetric complexes and each symmetry group is associated with unique structural and functional properties (Bergendahl and Marsh 2017; Goodsell and Olson 2000). The three most common symmetry groups in the human proteome are twofold symmetry (Schönflies notation, *C*_2_), higher-order cyclic symmetry (*C*_n>2_), and dihedral symmetry (*D*_n>1_). Members of the cyclic symmetry group frequently form pores in biological membranes (Forrest 2015), and because their subunits collectively contribute to function such as ion transport, they are prone to be afflicted by dominant mutations in their genes. Indeed, the fraction of AD genes is significantly higher in complexes with cyclic symmetry compared to those with twofold or dihedral symmetry (**Figure S1D**). By contrast, the dihedral symmetry group is enriched in metabolic enzymes (Bergendahl and Marsh 2017), which are often tolerant to heterozygous loss-of-function mutations owing to the law of diminishing returns in metabolic flux (Kacser and Burns 1981; Wright 1934).

We have previously shown that cotranslationally assembling members of the different symmetry groups tend to have larger interfaces (Badonyi and Marsh 2022), but they also differ in their level of cotranslational assembly, with members of the cyclic group being the least likely to cotranslationally assemble (**Figure S1E**). When we subset AD and AR homomers based upon their symmetry group, we find a relatively large variation in the level of cotranslational assembly (**Figure S1F**). A randomly selected cyclic complex with AD inheritance is four times less likely to cotranslationally assemble than an AR cyclic complex (OR = 0.24, p = 9.6 × 10^−5^), while a dihedral complex with AD inheritance is only half as likely (OR = 0.51, p = 0.045). Rare symmetries, such as helical and cubic, as well as asymmetric homomers, were grouped into the “other” category due to their relatively low representation in the human proteome. We did not detect a significant trend in this group, which may have to do with the heterogeneous properties of the complexes associated with those symmetries.

For example, torsin-A is a chaperone involved in synaptic vesicle recycling and it forms a helical complex with 8.5 subunits per turn (Demircioglu et al. 2019). The DN mutation E303del in torsin-A results in the deletion of a glutamate near the C terminus involved in early-onset torsion dystonia (Torres et al. 2004). Because complexes with helical symmetry are topologically open, they would undergo unlimited polymerisation under ideal conditions (Marsh and Teichmann 2015). Despite torsin-A having been detected to cotranslationally assemble (Bertolini et al. 2021), the large number of subunits and the nature of fibre assembly will likely render it susceptible to the DN effect, analogous to tubulin subunits that make up helical microtubules (Attard, Welburn, and Marsh 2022). This contrasts with comparably large complexes with a closed topology that could benefit from cotranslational assembly, such as the dihedral (D_39_) major vault protein, which has been observed to be readily assembled by the polyribosome (Mrazek et al. 2014).

Coiled coil motifs were found to be highly enriched among cotranslationally assembling proteins (Bertolini et al. 2021). Interestingly, we noticed that cotranslationally assembling homomers and repeated subunits are significantly enriched in alpha helices (effect size = 0.161, p = 1.5 × 10^−52^; Wilcoxon rank-sum test) and this enrichment is similar in effect after coiled coil motif containing proteins are removed from the data (effect size = 0.155, p = 9.6 × 10^−44^). We therefore controlled for alpha helix content as a potential confounder by binning the subunits into four quartiles (**Figure S1G**). The results indicate that the trend remains consistent across the bins, although lacks statistical significance for the bin with the lowest amount of helix content.

Bertolini *et al*. have provided confidence-based classification for the cotranslationally assembling protein candidates (Bertolini et al. 2021). While the exclusive use of high confidence proteins is prohibitive to the analysis, as they comprise only one-fifth of all detected proteins, we controlled for its potential confounding effect by splitting the subunits into high and low confidence groups (**Figure S1H**). The high confidence group excludes all low confidence proteins, and similarly, the low confidence group excludes high confidence proteins altogether. We found that the level of cotranslational assembly is significantly lower in AD relative to AR subunits in both high (OR = 0.66, p = 7.7 × 10^−3^) and low confidence groups (OR = 0.55, p = 5 × 10^−10^). Because, by design, high confidence candidates are exclusively cytoplasmic or nuclear proteins, one possible explanation to why the trend is stronger in the low confidence subset is that they are enriched in membrane-bound complexes that commonly adopt a cyclic symmetry, which demonstrates the strongest buffering capacity among the main symmetry groups (**Figure S1F**).

We also analysed the level of cotranslational assembly in the subset of AD genes that are also associated with AR disorders (AD/AR). In **Figure 2F**, we show that AD/AR genes of homomers and repeated subunit heteromers are more strongly depleted in cotranslational assembly relative to AR (OR = 0.49, p = 1.4 × 10^−7^; Holm-Bonferroni corrected hypergeometric test) than the AD group (OR = 0.61, p = 3.7 × 10^−7^). Because, as mentioned earlier, dominant mutations in mixed AD/AR subunits often phenocopy the recessive disorder, they are thought to be enriched in assembly-mediated DN and dominant-positive effects, which are more likely to come about in the absence of cotranslational assembly.

Finally, we considered genetic dominance of mutations in genes located on autosomes and on the X chromosome. Genes on autosomes are normally present in two copies, one allele on each chromosome, corresponding to the diploid homo-or heterozygous states. Genes on the X chromosome are hemizygous in males, meaning that only one allele is present at all times. In females, the second allele is generally silenced with the exception of a subset of genes that escape the inactivation process in a tissue-specific manner (Tukiainen et al. 2017). Consequently, cotranslational assembly should have no capacity to buffer X-linked dominance, since there is not a wild-type allele whose transcript could ameliorate the phenotype. In accordance with this idea, genes of homomers and repeated subunits that cause disease via an X-linked dominant pattern do not appear to have lower levels of cotranslational assembly than their X-linked recessive counterparts (**Figure S1I**, OR = 1.06, p = 0.639).

### Subunits with dominant-negative disease mutations are less likely to cotranslationally assemble than subunits with heterozygous loss-of-function mutations

Given that the fraction of cotranslationally assembling homomers and repeated subunits is lower in AD relative to AR genes, we sought to identify which molecular mechanism in the AD inheritance mode is responsible for the trend. Dominant molecular mechanisms by which mutations cause disease at the protein-level can be broadly classified into three groups: heterozygous loss of function (LOF, or haploinsufficiency), gain of function (GOF), and dominant negative (DN). Because DN mutations adversely affect the function of the wild type allele, we hypothesised that subunits with DN disease mutations should be depleted in cotranslational assembly relative to genes with LOF mutations that inactivate one allele without affecting the wild type. This is because cotranslational assembly should reduce the mixing that occurs between the wild-type and mutant subunits, which in turn should make it less likely that the assembled complex will exhibit a disproportionate loss-of-function, i.e. a DN phenotype.

To start with, we classified 66% of AD genes (n = 1,185; available in **Table S1**) into LOF, GOF, and DN molecular mechanisms using text-mining approaches and manual curation of the corresponding evidence (detailed in **Methods**). To ascertain that the gene sets are suitable for analysis, we have looked at properties known to be associated with the different molecular mechanisms. These include the change in Gibbs free energy (ΔΔG) upon pathogenic mutations and the extent to which the mutations cluster in 3D space (Gerasimavicius, Livesey, and Marsh 2022). It was first observed in a subset of membrane proteins that DN mutations tend to have low (predicted) ΔΔG values (McEntagart et al. 2016). This may be explained by the idea that non-LOF mutations should generally permit at least the partial folding of the subunit and so they tend to have a mild impact on the structure. Conversely, LOF mutations inactivate the protein, which frequently happens when protein interior residues are involved whose mutations are considerably more structurally damaging. In agreement with this, we found that disease mutations in homomers and repeated subunits of the DN class have significantly lower predicted ΔΔG compared to LOF subunits (**Figure S2A**, p = 1.3 × 10^−16^; Wilcoxon rank-sum test). In terms of 3D clustering, non-LOF mutations are often concentrated in specific regions of a protein (Lelieveld et al. 2017), such as interfaces and functional sites, while LOF mutations tend be uniformly distributed across the structure. Consistent with this, disease mutations of DN homomers and repeated subunits in our dataset exhibit a higher degree of 3D clustering than LOF subunits (**Figure S2B**, p = 4.7 × 10^−4^; Wilcoxon rank-sum test), with a strong tendency to be enriched at homomeric interfaces (**Figure S2C**, p = 1.3 × 10^−18^; hypergeometric test).

We next directly addressed our hypothesis by calculating the fraction of cotranslationally assembling subunits in each molecular mechanism group, shown in **Figure 3A**. In agreement with the hypothesis, we found the fraction to be markedly lower among DN (20.9%) compared to LOF subunits (32.4%, OR = 0.55, p = 1.5 × 10^−3^, hypergeometric test) and AR subunits (35.6%, OR = 0.49, p = 6.9 × 10^−8^). At the molecular level, AR and heterozygous LOF mutations are very similar in their effect. Recessive disorders are almost always due to biallelic (homozygous or compound heterozygous) loss of function, with a few rare examples of biallelic gain of function (Cavaco et al. 2018; Drutman et al.

**Figure 3.**
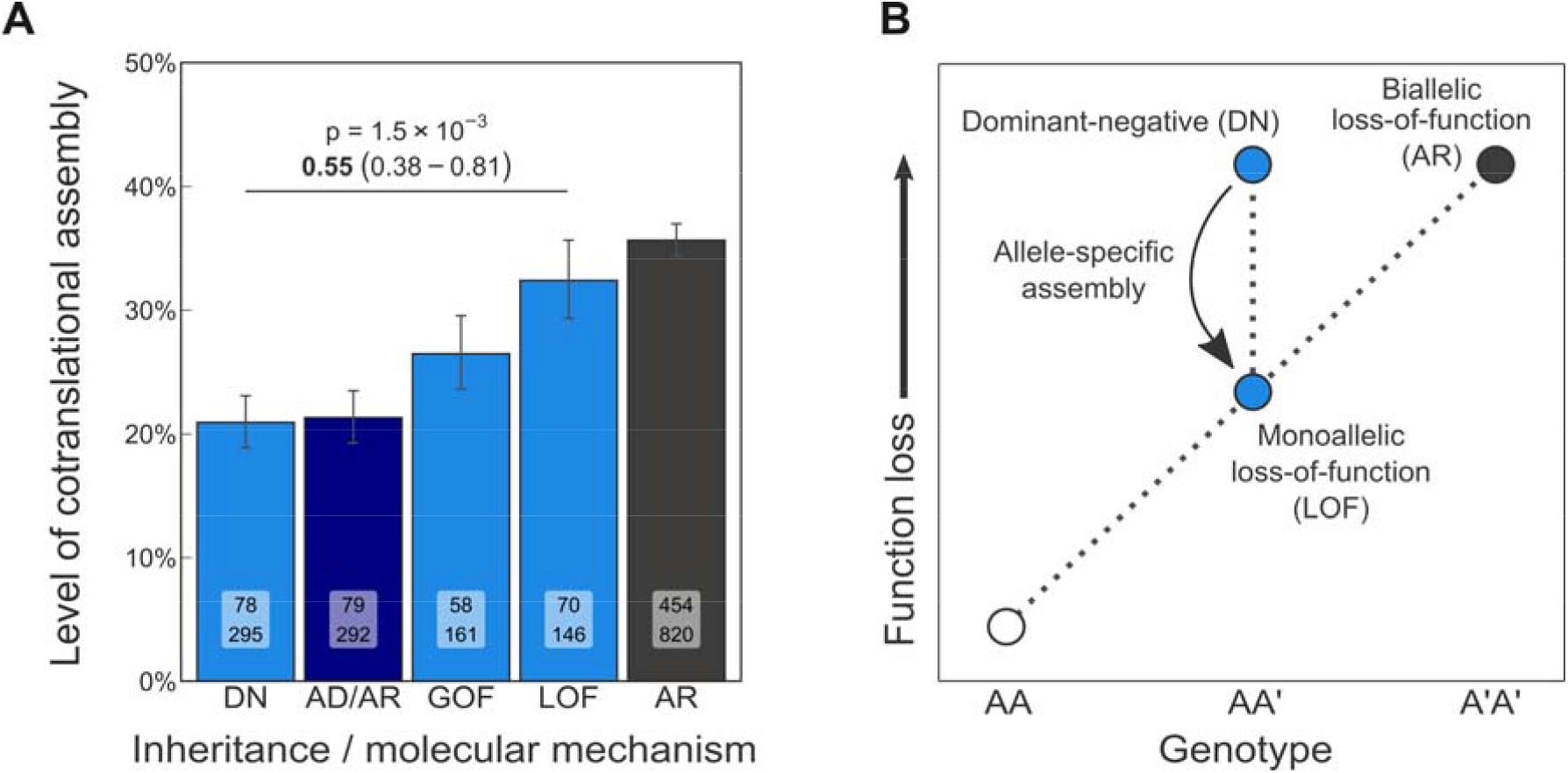
Subunits with dominant-negative disease mutations are less likely to cotranslationally assemble than subunits with heterozygous loss-of-function mutations (**A**) Level of cotranslational assembly in homomers and repeated subunit heteromers according to autosomal recessive inheritance (AR), mixed autosomal dominant / autosomal recessive inheritance (AD/AR), and dominant molecular disease mechanisms: loss of function (LOF), gain of function (GOF), and dominant negative (DN). The hypergeometric p-value and the odds ratio (OR, in bold) and its confidence interval of the DN vs LOF comparison is shown. (**B**) Genotype-function loss landscape diagram. Allele-specific assembly of homomers and repeated subunit heteromers may alleviate the effect of a monoallelic (heterozygous) LOF mutation by reducing the mixing between the product of the wild-type and the mutant genes.

2019; Wishner et al. 1975). Altogether, these results indicate that the level of cotranslational assembly in subunits with monoallelic and biallelic LOF mutations is similar, but subunits with DN mutations are observed to cotranslationally assemble less frequently. Therefore, allele-specific protein complex assembly may prevent some mutations from the clinical manifestation of a phenotype caused by certain heterozygous variants (**Figure 3B**).

Further indication that mutations in genes with both dominant and recessive inheritance are strongly associated with assembly mediated effects is borne out in the molecular mechanism classifications, with the DN class having the highest fraction of genes with mixed inheritance (36.6%) followed by GOF (29.2%) and then LOF (17.8%). When LOF genes are removed from the AD/AR set to allow for a non-redundant comparison, the fraction of AD/AR genes that cotranslationally assemble is 18.4%, which is significantly less than the 32.4% measured in the LOF class (OR = 0.47, p = 2.8 × 10^−4^; hypergeometric test). Furthermore, we observe the GOF class to be depleted in cotranslational assembly relative to LOF, although not to a statistically significant extent (26.5% vs 32.4%, p = 0.11). We speculate that this observation is driven by the enrichment of the GOF class in the assembly-mediated dominant-positive effect, which is subject to genetic buffering by allele-specific assembly. We also noticed that GOF homomers more frequently assemble into cyclic complexes relative to the LOF class (21.3% vs 6.4%, p = 3.9 × 10^−3^; Fisher’s exact test; **Figure S2D**), the symmetry group with the lowest fraction of cotranslationally assembling members (18.4% compared to 27.3% in twofold homodimers and 31.2% in dihedral complexes; **Figure S1E**).

For the above reason, we explored the potential confounding effect of structural symmetry, and found prominent symmetry group preferences among the molecular mechanisms (**Figure S2D**). DN homomers, similar to the GOF homomers, are enriched in the cyclic symmetry relative to LOF (20.4% vs 6.4%, p = 4 × 10^−3^; Fisher’s exact test). LOF homomers frequently form *C*_2_ symmetric dimers, although their sample size does not allow for a statistically robust conclusion. Moreover, AR homomers are overrepresented in the dihedral symmetry group relative to DN homomers (12.6% vs 4.8%, p = 1.3 × 10^−3^; Fisher’s exact test). We suspect that these symmetry group compositions reflect biases in protein function. The inheritance and molecular mechanism of disease strongly correlate with protein function. For example, disorders caused by genes encoding enzymes are primarily recessive (Jimenez-Sanchez, Childs, and Valle 2001), genes encoding transcription factors are more likely to be haploinsufficient (Seidman and Seidman 2002), and those of membrane channels commonly give rise to DN and GOF mutations (Celesia 2001). These admittedly broad functional classes have been shown to be associated with structural properties, with metabolic enzymes being enriched in dihedral symmetry, transcription factors in twofold symmetry, and membrane channels in cyclic symmetry (Bergendahl and Marsh 2017; Forrest 2015; Goodsell and Olson 2000). We used protein functional classification to investigate if the aforementioned functions are reflected in the inheritance and molecular mechanism classes, which supported the above assumptions (**Figure S2E**). Metabolic enzymes are overrepresented in AR subunits (AR vs all other, OR = 4.27, p = 3.9 × 10^−40^; Holm-Bonferroni corrected Fisher’s exact test), membrane transporters among GOF and DN subunits (OR = 2.96, p = 4.8 × 10^−9^ and OR = 1.99, p = 4.8 × 10^−6^, respectively), and LOF subunits are 6.9-fold more likely to function as transcription factors than a subunit sampled randomly from the disease gene pool (OR = 6.89, p = 2.15 × 10^−19^).

When homomers in the different molecular mechanism classes are grouped by their symmetry, the level of cotranslational assembly is consistently lower among DN subunits and those with mixed AD/AR inheritance than LOF or AR subunits (**Figure S2F**). The only exception is the cyclic symmetry in the LOF class, where it is difficult to reliably estimate the cotranslational assembly rate because no cotranslationally assembling member was identified. Furthermore, we split the analysis into homomers and repeated subunits to examine if the trend seen in **Figure 3A** holds up for repeated subunits within the dominant molecular mechanisms. We show in **Figure S2G** that strictly homomers with DN mutations are still more strongly depleted in cotranslational assembly than LOF homomers (OR = 0.44, p = 1.1 × 10^−3^; hypergeometric test) and that repeated subunits with AD/AR inheritance are also depleted relative to AR (OR = 0.47, p = 5.7 × 10^−4^). These results demonstrate that the genetic buffering capacity of cotranslational assembly, despite evident differences in structural symmetry, is neither confounded by symmetry nor is it exclusive to homomers but includes also repeated subunits of heteromeric complexes.

### Interfaces of homodimers with dominant-negative disease mutations are C-terminally shifted

How can we explain the observation that subunits with DN disease mutations are less likely to cotranslationally assemble? As previously discussed, two potential confounding variables are protein abundance and length. However, we did not find significant differences in the abundance and length distributions of homomers and repeated subunits in the dominant molecular mechanism classes that could explain the trend (Dunn’s test of multiple comparisons with Holm-Bonferroni correction, **Figure S3A-B**). This suggests that there must be other properties of DN subunits that make them less likely to assemble on the ribosome. There are three reported structural features that show correspondence with cotranslational assembly: high alpha helix content, a large interface area, and an N-terminally localised interface (Bertolini et al. 2021; Kamenova et al. 2019; Natan et al. 2018; Shiber et al. 2018). We reasoned that the difference should reside in these structural features given the inherently mechanistic nature of cotranslational assembly.

We first investigated the propensity of homomeric and repeated subunits to form alpha helices. Surprisingly, we found that LOF subunits have fewer alpha helical secondary structure elements than DN subunits (26.7% vs 33.9%, p = 3.9 × 10^−5^, Dunn’s test of multiple comparisons with Holm-Bonferroni correction, **Figure S3C**) or AR subunits (33.3%, p = 2.8 × 10^−7^). Although helix propensity correlates with cotranslational assembly, it is possible that there are stronger functional constraints that determine whether or not a complex forms on the ribosome. We thought that the low helix content in the LOF group may be indicative of high protein disorder, as disorder is naturally anticorrelated with structured regions in proteins. To investigate this, we took advantage of the structure-averaged pLDDT (predicted local-distance difference test) values from the AlphaFold predicted human protein models, which are highly predictive of intrinsic disorder (Akdel et al. 2022; Tunyasuvunakool et al.

2021). As expected, LOF homomers and heteromers exhibit the lowest median pLDDT value (median = 75.7, **Figure S3D**), significantly lower than DN subunits (median = 77, p = 1.6 × 10^−5^, Dunn’s test of multiple comparisons with Holm-Bonferroni correction) and AR subunits (median = 83.3, p = 1.4 × 10^−6^). Therefore, the structures of LOF subunits are likely to contain a high fraction of disorder. This conclusion is consistent with our earlier result showing that LOF subunits are almost seven-fold more likely to be transcription factors than other disease genes, because transcription factors are known to be strongly associated with intrinsic disorder (Liu et al. 2006).

Transcription factor subunits may benefit from cotranslational assembly as a means of preventing their toxicity and fine-tuning the dosage and timing of their expression during developmental processes and the cell cycle (Schwarz and Beck 2019).

We next assessed the influence of subunit interface area in homomeric subunits, which we have shown to have a strong relationship with cotranslational assembly (Badonyi and Marsh 2022).

Homomeric complexes with larger subunit:subunit contact areas are more hydrophobic and therefore experience a stronger drive to assemble as soon as their interfaces become exposed on the ribosome. Due to the unique structural and functional characteristics associated with homomeric symmetry, we split the subunit interface areas based upon their symmetry group. The analysis revealed that subunits associated with DN disease mutations do not have smaller interfaces than most other disease associated subunits that would make them less likely to cotranslationally assemble (**Figure S3E**). In fact, interface areas of LOF homomers are significantly smaller than those of the DN subunits in the main symmetry groups (**Figure S3E**). On one hand, this result is consistent with the enrichment of pathogenic mutations at interfaces of DN subunits (**Figure S2C**), which, assuming a random mutation model, would be less likely to occur if the interfaces were small. On the other hand, GOF, DN, as well as AD/AR subunits may have larger interface areas simply because it reflects the biological significance of their interfaces, since their molecular mechanisms are dependent upon complex assembly. Conversely, subunits in the LOF group may have a higher fraction of crystallographic interfaces, which tend to be smaller (Prasad Bahadur et al. 2004).

The third structural property we investigated is the relative interface location (Badonyi and Marsh 2022). N-terminal regions of proteins are more likely to be involved in cotranslational interactions because of vectorial synthesis on the ribosome, an intuitive assumption supported by experimental observations (Bertolini et al. 2021; Kamenova et al. 2019; Natan et al. 2018; Shiber et al. 2018). We hypothesised that interfaces of homomeric subunits with DN disease mutations should be C-terminally shifted, which would strongly decrease their tendency to cotranslationally assemble. We first calculated the relative interface location for all homomeric subunits and the average interface location of the symmetry groups (**Figure S3F**). Based upon the difference between the symmetry average and the subunits’ relative interface location, we found that the interfaces of twofold symmetric dimers with DN disease mutations are significantly more C-terminal than what is expected from the symmetry group (p = 0.025; Holm-Bonferroni corrected Wilcoxon rank-sum test against basemean). To quantify this difference, we resampled the dataset of twofold homodimers with replacement stratified by molecular mechanisms and calculated confidence intervals. **Figure 4A-B** show the raw relative interface location values as well as the bootstrap distribution of the relative interface shift, i.e. distance from the symmetry mean. The relative interface shift can be interpreted as a percentage, where +5% indicates that the subunit’s observed interface is C-terminal to its expected value by 5% of the protein’s length. According to this, subunits of homodimers with DN disease mutations are 6% more C-terminal than the symmetry group mean (p = 7 × 10^−3^, resampling p-value).

**Figure 4.**
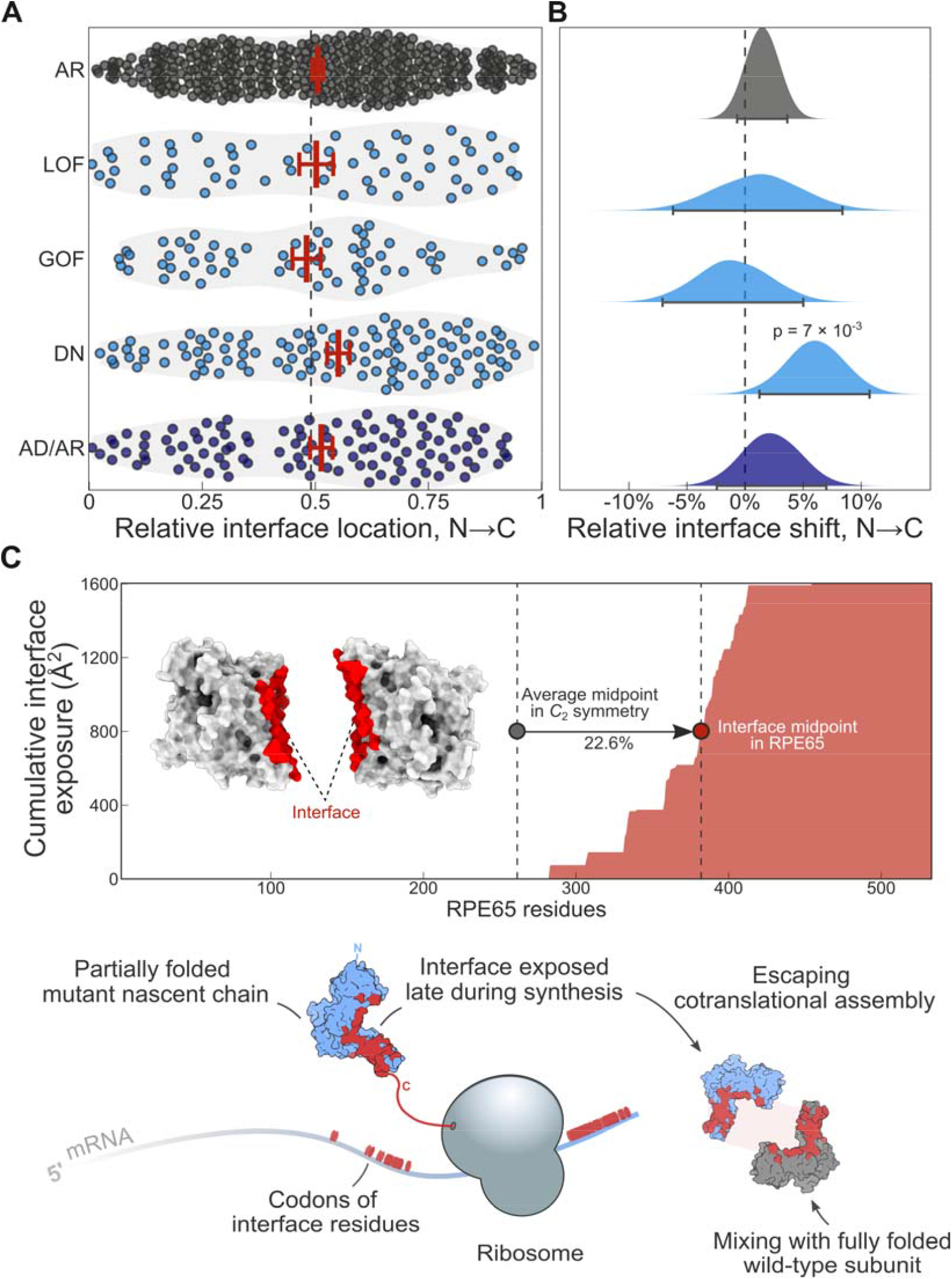
Interfaces of homodimers with dominant-negative disease mutations are C-terminally shifted (**A**) Relative interface location of *C*_2_ symmetric homodimers according to autosomal recessive inheritance (AR), mixed autosomal dominant / autosomal recessive inheritance (AD/AR), and dominant molecular disease mechanisms: loss of function (LOF), gain of function (GOF), and dominant negative (GOF). The dashed line is at 0.49, which is the average interface location of the symmetry group. Crossbars are mean ± SEM. (**B**) Bootstrap distributions of the difference between the symmetry average relative interface location and the observed value for *C*_2_ symmetric homodimers in the different classes. Error bars represent 95% percentile confidence intervals and the p-value was calculated from the resamples. (**C**) Top: Cumulative interface exposure of the enzyme RPE65 during the translation process. Half of its final interface area (1,604 Å^2^) is reached 22.6% later than what is expected from the symmetry group. Bottom: Mechanistic interpretation of C-terminally shifted interfaces in DN subunits with *C*_2_ symmetry.

We illustrate the mechanistic interpretation of this finding in **Figure 4C**. The retinoid isomerohydrolase (RPE65) is an enzyme involved in the conversion of all-trans-retinyl esters into 11-cis-retinol, which is essential for phototransduction in the retinal pigment epithelium (Kiser et al. 2015). Both autosomal recessive and dominant mutations in *RPE65* have been associated with retinitis pigmentosa, which makes heterozygous mutations in the gene less likely to act via simple LOF mechanisms. The DN mutation D477G was found to exert a DN effect on the wild-type and lead to delayed chromophore regeneration (Bowne et al. 2011). Interestingly, the interface of RPE65 during translation is exposed 22.6% later than what would be expected from an average *C*_2_ dimer, which is a condition that does not favour assembly on the ribosome. Consistently, RPE65 was not detected to cotranslationally assemble by disome selective ribosome profiling (Bertolini et al. 2021). These observations support a model whereby subunits that expose their interface residues late in the translation process are less likely to cotranslationally assemble and as a result, are more likely to be associated with DN mutations.

### Training and assessing a classifier for the prioritisation of genes with alternative molecular mechanisms

There have been important studies aimed at understanding the properties of haploinsufficient genes and their prediction (Dang et al. 2008; Huang et al. 2010; Karczewski et al. 2020; MacArthur et al. 2012; Petrovski et al. 2015; Shihab et al. 2017; Steinberg et al. 2015), but comparatively little effort has been channelled into exploring the characteristics of genes that give rise to dominant disorders in a manner not explained by simple LOF. Recently, we have shown that variants in genes associated with non-LOF mechanisms are predicted substantially poorer by state-of-the-art variant effect predictors (Gerasimavicius, Livesey, and Marsh 2022). This observation emphasises the importance of considering alternative molecular mechanisms in our collective attempt to annotate the human pathogenic variation. While the text-mining strategy we utilised here was able to provide likely molecular mechanism assignments for 1,185 dominant disease genes, many others remain unknown. Moreover, for as-of-yet undiscovered disease genes, there is a strong possibility that we could miss pathogenic mutations due to their association with DN or GOF mechanisms and thus worsen computational predictions. To facilitate future variant-level molecular mechanism prediction and aid clinical geneticists evaluate the potential of novel inherited or *de novo* mutations to inflict a non-LOF consequence on the protein, we built a classifier with the goal of identifying genes most likely to be associated with non-LOF genes over those that are primarily associated with LOF mechanisms.

We first reviewed the literature to identify properties of LOF genes and then trained a logistic regression model with lasso penalty using a range of diverse features, including cotranslational assembly (Bertolini et al. 2021), the functional and structural determinants investigated in this study, as well as population-level mutational constraints (Karczewski et al. 2020), evolutionary-, sequence-and interaction network-based properties (Huang et al. 2010; Shihab et al. 2017; Steinberg et al. 2015), and experimental data (Uhlen et al. 2010; M. Wang et al. 2015) (detailed in **Methods**). Measured on the test set, the classifier achieves a receiver operating characteristics area under the curve of 0.74 (**Figure 5A**), an F_1_ score of 0.8, and a Matthews correlation coefficient of 0.24 (Chicco, Tötsch, and Jurman 2021) (detailed performance profile in **Figure S4**). Cotranslational assembly was found to be a discriminating feature in the model, ranking 12^th^ out of 30 features and being roughly one-fourth as important as the top predictor, which is the ratio of nonsynonymous-to-synonymous substitutions (dN/dS) in the coding sequence of human relative to macaque genes (Huang et al. 2010) (**Figure 5B**). Notably, the second and third most important predictors in the model are transporter/channel function and the number of paralogues of the gene. It has been observed before that haplosufficient genes have higher average sequence identity to the closest paralogue than LOF genes (Huang et al. 2010), suggesting functional compensation by closely related proteins. It is possible that due to this functional redundancy, autosomal dominant genes with a high number of paralogues are simply more likely to be associated with non-LOF mechanisms. For example, ion channel genes are known to have undergone multiple gene duplication events (Liebeskind, Hillis, and Zakon 2015), which is consistent with their enrichment among DN and GOF subunits (**Figure S2D/E**). In cyclic complexes with more than one unique subunit, paralogous copies typically sequester in the same complex (Mallik, Tawfik, and Levy 2022), suggesting that information on paralogues is a valuable proxy for non-LOF mechanisms in homomers as well as in heteromers.

**Figure 5.**
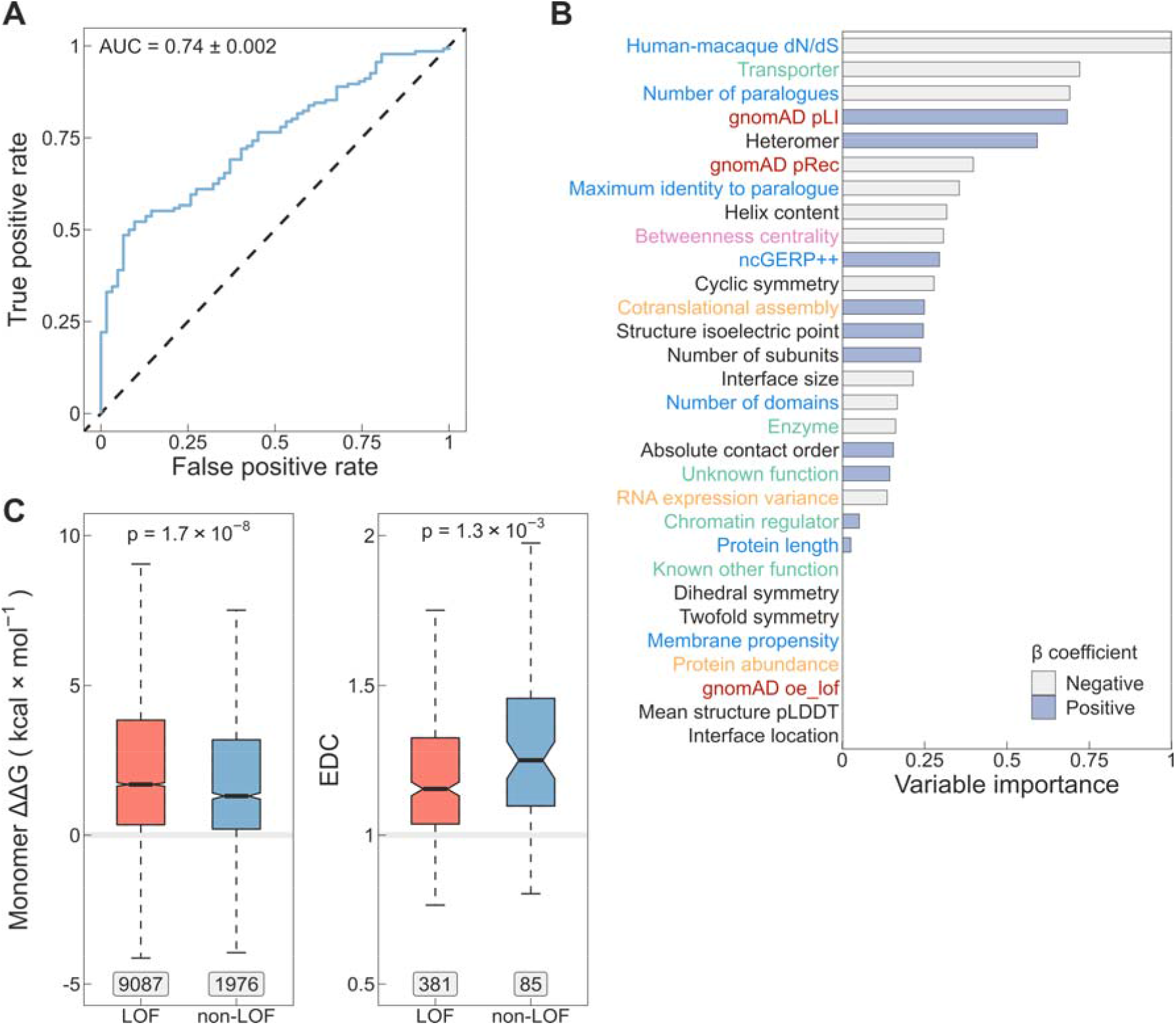
A computational model for identifying genes most likely to be associated with non-LOF molecular mechanisms (**A**) Receiver operating characteristic (ROC) curve of the lasso regression model measured on the test set. AUC ± bootstrap (n = 1,000) standard error is shown. (**B**) Variable importance calculated as the absolute values of the β coefficients scaled to the [0,1] interval. The y-axis labels are coloured according to the type of the variable: sequence-derived or evolutionary variables (blue), functional annotations (green), mutational constraint metrics (red), structural properties (black), interaction network-based property (pink), and experimental data (orange). Bars are coloured based upon the sign of β. (**C**) Differences in Gibbs free energy change (left) and extent of disease clustering (EDC, right) of pathogenic mutations between genes predicted to be non-LOF versus all other genes at threshold T2 Genes that were used for training the model as well as known autosomal recessive genes were excluded. Boxes denote data within 25^th^ and 75^th^ percentiles, the middle line represents the median, the notch contains the 95% confidence interval of the median, and the whiskers extend from the upper and lower quartiles to a distance of 1.5 times the interquartile range. Labels indicate the number of variants (ΔΔG) or the number of genes (EDC) in the groups. The p-values were calculated with Wilcoxon rank-sum tests.

We derived two probability thresholds (**Figure S4E**). The threshold of p = 0.82 (T1) was selected on the basis of the maximum value of Youden’s J statistic (Youden 1950) (test set confusion matrix: 68/68 [50%] non-LOF vs 5/57 [8%] LOF). A second threshold of p = 0.92 (T2) was chosen as the value at which the specificity of the model reaches 100%, i.e. no ground truth LOF genes are classified as non-LOF at the cost of classifying more ground truth non-LOF genes as LOF (29/107 [21%] non-LOF vs 0/62 [0%] LOF). We provide predictions for 9,051 proteins covering ∼44% of the proteome (available in **Table S2**) that have at least partial structures in the PDB. Of these, 880 (9.7%) are above T2 and 3,315 (36.6%) are above T1. Of the latter, 2,840 have no dominant disease association recorded in OMIM.

As an unbiased approach to assess the model, we analysed the ΔΔG of pathogenic mutations and their extent of disease clustering (EDC) after removing genes used for training, and AR genes, whose biallelic LOF mutations would bias the trend. In **Figure 5C** we show the result of this analysis at threshold T2, demonstrating that missense mutations in predicted non-LOF genes exhibit a milder impact on protein structure (Gerasimavicius, Livesey, and Marsh 2022; McEntagart et al. 2016). Moreover, pathogenic variants in predicted non-LOF genes show strong 3D clustering in their respective protein structures, consistent with previous observations (Gerasimavicius, Livesey, and Marsh 2022; Lelieveld et al. 2017). In **Figure S4F/G**, we provide further support that, between thresholds T1 and T2, both ΔΔG and EDC exhibit the expected trend with an increasing effect size.

## Discussion

Cotranslational assembly of homomers is thought to result in complexes whose subunits originate from the same allele (Bertolini et al. 2021; Gilmore et al. 1996; Mrazek et al. 2014; Redick and Schwarzbauer 1995). A possible consequence of this mechanism is that cotranslationally assembling subunits harbouring pathogenic heterozygous mutations may be sequestered into half of the protein complex pool rather than mixing with the wild-type and inflicting functionally harmful effects. By comparing the fraction of cotranslationally assembling subunits associated with Mendelian diseases, we showed that genes of homomers and repeated subunits inherited in an AD pattern are significantly depleted in this mode of assembly compared to AR genes. Moreover, we found that among AD genes of homomers those that exert a DN effect are the least likely to cotranslationally assemble compared to other protein-level molecular mechanisms of disease, but importantly, to those that predominantly harbour heterozygous LOF mutations. Our results therefore reveal a previously hypothesised genetic buffering mechanism (Natan et al., 2017; Perica et al., 2012), whereby complexes undergoing cotranslational assembly are to some extent protected from the deleterious consequences of DN mutations.

We observe AR complexes to consistently have high levels of cotranslational assembly regardless of their structural symmetry. It was first proposed by Wright (Wright 1929) and Haldane (Haldane 1930), whose ideas were developed further by Hurst and Randerson (Hurst and Randerson 2000), that recessivity is a consequence of selection for larger amounts of protein, because the high abundance of enzymes is a “safety factor” (Kacser and Burns 1981) that increases their robustness to dominant mutations. It is possible that the high abundance and the structural properties of metabolic enzymes, such as their enrichment in the dihedral symmetry (Bergendahl and Marsh 2017), necessarily lead to frequent cotranslational assembly events, representing an additional safety factor against the deleteriousness of dominant, especially DN mutations. Although the evolution of protein oligomeric state can arise from nonadaptive processes (Hochberg et al. 2020; Johnston et al. 2022; Lynch 2013), it is not implausible that biological phenomena such as this impose weak selection.

Our results hint at the extraordinary regulation of protein complex assembly within cells. Allele-specific assembly in homomers may emerge from the inherent colocalisation of their nascent chains, although certain protein structural features appear to predispose subunits to the process. Interestingly, we observed repeated subunits of heteromeric complexes to also exhibit genetic buffering by cotranslational assembly. According to one hypothesis, subunits may combine information in their mRNAs and protein sequences to increase the efficiency of assembly mediated by RNA-binding proteins (Chen and Mayr 2022; Fabián Morales-Polanco et al. 2022). A range of membraneless compartments have been put forward as putative sites of intense protein complex assembly under physiological conditions, including TIS granules (Ma and Mayr 2018), assemblysomes (Panasenko et al. 2019), and translation factories (Fabian Morales-Polanco et al. 2021), which may well represent the same type of condensates (reviewed in Fabián Morales-Polanco et al. 2022).

Across diverse proteomes, interface contacts of homomers are enriched towards the C terminus, which is thought to be the product of evolutionary pressure on folding to happen before assembly (Natan et al. 2018). By contrast, N-terminal protein interfaces have been found to favour cotranslational assembly (Badonyi and Marsh 2022; Bertolini et al. 2021; Kamenova et al. 2019; Natan et al. 2018; Shiber et al. 2018). Interestingly, our structural analysis suggests that interfaces of homodimers with DN disease mutations are significantly shifted towards the C-terminus relative to what is expected from their symmetry group. As a possible consequence, their interfaces become exposed in nascent polypeptides relatively late during translation, strongly reducing their likelihood of cotranslational assembly. This exciting observation represents a survivorship bias, so that we tend to observe subunits cause disease via a DN mechanism when they “escape” cotranslational assembly and subsequently co-assemble with wild-type subunits.

Ongoing efforts to develop variant effect predictors focusing on the molecular consequences of protein-coding variants should consider whether the subunit cotranslationally assembles or, providing that structural data on the complex is available, the properties of interfaces in order to prioritise those with a possible DN effect. As demonstrated in this study, a substantial fraction of subunits with mutations inherited in an AD pattern display a DN phenotype, likely including many of those that have not yet been characterised well enough to be assigned to one of the protein-level dominant molecular mechanism classes. Moreover, current clinical sequencing pipelines frequently identify inherited and recurrent *de novo* heterozygous variants in genes that have historically been associated with recessive disorders, which are often ranked lower under the assumption that they would not be pathogenic in a heterozygous state (Birgmeier et al. 2020; Eilbeck, Quinlan, and Yandell 2017; Paila et al. 2013; G. T. Wang, Peng, and Leal 2014; K. Wang, Li, and Hakonarson 2010; Zemojtel et al. 2014). However, a DN effect is possible if the gene encodes a homomer or repeated subunit heteromer (Backwell and Marsh 2022), and especially if its (sub)complex assembles post-translationally. Here, we provide predictions of non-LOF genes covering almost half of the human proteome to expedite the discovery of new variants associated with alternative molecular disease mechanisms.

Finally, our results shine light onto the fascinating connection between inheritance, which determines the genetic traits of an individual, and protein complex assembly, which takes place only after the genetic information has been decoded. Future work will be required to directly measure the effect of allele-specific assembly on the mixing of wild-type and mutant subunits in vivo.

## Methods

### Structural data

We searched the Protein Data Bank (PDB) (Berman et al. 2000) on 2021-02-18 for all polypeptide chains >50 amino acids and >90% sequence identity to human canonical sequences in the UniProt proteome UP000005640. For genes that map to multiple chains, we selected a single chain ranking by sequence identity, the number of unique subunits in the complex, and by the number of atoms present in the chain. In every case we used the first biological assembly and its symmetry assignment was taken from the PDB. The interface area was calculated at residue-level between all pairs of subunits with AREAIMOL from the CCP4 suite (Winn et al., 2011), using a probe radius of 1.4 Å. The interface was defined as the difference between the solvent accessible surface area of the subunit in isolation and within the context of the full complex. Subunits with interfaces >400 Å^2^ were considered for analysis to exclude potentially crystallographic interfaces.

We extended the PDB dataset with homology models of human homomeric complexes in the SWISS-MODEL repository (UniProt release 2022_02) (Bienert et al., 2017; Waterhouse et al., 2018). Models based on isoform sequences were excluded. The software AnAnaS (Pagès et al., 2018; Pagès &Grudinin, 2018) was run on default settings to determine the number of subunits and the symmetry group of the complexes. In rare cases when symmetry was not detected, we assigned the symmetry group of the PDB template used to model the complex. If a protein was found in multiple homology models, we selected the one with the largest number of subunits followed by the length of the modelled chain. The interface area was calculated at residue-level between all pairs of subunits with FreeSASA 2.0.3 (Mitternacht, 2016), using a probe radius of 1.4 Å. We performed pairwise alignments between the modelled chain and the paired UniProt sequence to confirm residue correspondence to the canonical sequence, because any mismatch in residue numbering could influence the relative interface location metric. Similarly to the PDB structure data, only subunits with interfaces >400 Å^2^ were included in the analyses. When the SWISS-MODEL dataset was pooled with the PDB dataset, we prioritised the homomeric subunit with the larger interface area.

### Relative interface location

The relative interface location is a value between 0 and 1 indicating the location of the interface relative to the protein termini (N=0 and C=1), and it was calculated as previously described (Badonyi and Marsh 2022). To ensure the analysis is not biased by homologous proteins, we generated a distance matrix based on the sequences of the chains from the structures using Clustal Omega version 1.2.4 (Sievers et al. 2011). The distances were converted to per cent identities and the matrix was filtered to below 50% using a redundancy filtering algorithm (Peeters et al. 2019). Only those structures were included in the analysis that passed the homology cutoff.

### FoldX free energy calculation

FoldX 5.0 (Delgado et al. 2019) was used to calculate the change in Gibbs free energy of ClinVar (Landrum et al., 2018) missense mutations in AlphaFold predicted structures of human proteins (Tunyasuvunakool et al. 2021). The “RepairPDB” command was first run to minimise structures followed by the “BuildModel” command on the repaired structures. The final Gibbs free energy change was calculated as the average of ten replicates, and in subsequent analyses residues with pLDDT < 50, which are predicted to be disordered in solution (Akdel et al. 2022), were excluded.

### 3D clustering of missense pathogenic mutations

The extent of disease clustering (EDC) metric expresses the proximity of every disease non-associated protein residue to a known disease-associated residue, and it was calculated as previously described (Gerasimavicius et al., 2022) from AlphaFold predicted structures. Briefly, for each residue with pLDDT > 50, we calculated the alpha carbon distance to all other residues with a known ClinVar disease mutation, selecting the shortest distance. The final metric is derived as the ratio of the common logarithm of non-disease and disease average distances. Values ≤1 indicate that the mutations are dispersed and those >1 suggest a degree of spatial clustering. Only proteins with at least 3 pathogenic or likely pathogenic missense mutations in ClinVar were included.

### Alpha helix content and mean pLDDT

The percentage of alpha helix residues was calculated from the AlphaFold predicted structures of human proteins (Tunyasuvunakool et al. 2021) using DSSP version 2.2.1 (Kabsch and Sander 1983). The per-residue pLDDT (predicted local-distance difference test) metric in the temperature factor field averaged over the structures to yield a per-protein confidence score.

### Gene-level inheritance patterns

Gene-disease inheritance relationships were obtained from OMIM (Amberger et al. 2015). Gene-specific XML files were retrieved via the OMIM application programming interface in 4 batches over consecutive days ending on 2022-07-07. Inheritances were extracted from the “phenotypeInheritance” node of each XML file.

### Gene set of homomers and repeated subunits

We extended the gene set of homomers identified by our structural mapping pipeline with genes that have non-structural evidence to form homo-oligomers or are present in >1 copy in a complex. For homomers, we used UniProt (Bateman et al., 2021), EMBL-EBI ComplexPortal (Meldal et al., 2019), CORUM (Giurgiu et al., 2019), the OmniPath database (Dénes et al. 2021), as well as single spanning membrane homodimers from the Membranome 3.0 database (Lomize et al. 2022). For repeated subunits, we extracted protein chains that appear in multiple copies in the biological units of complexes in the PDB (Marsh et al. 2015), and included proteins that have a stoichiometry >1 in the OmniPath database. Homomers were removed from the repeated subunit list to create a non-redundant dataset.

### Gene-level classification of dominant molecular mechanisms

We classified autosomal dominant genes into molecular disease mechanisms via text-mining PubMed (Sayers et al. 2021) titles and abstracts and OMIM XML gene entries. We searched PubMed using the keywords “dominant negative” for the dominant-negative (DN) mechanism, “gain of function” OR “activating mutation” for the gain-of-function (GOF) mechanism, and “haploinsufficiency” OR “haploinsufficient” OR “dosage sensitivity” OR “dosage sensitive” OR “heterozygous loss of function” for the loss-of-function (LOF) mechanism. The same workflow was applied to OMIM entries. The resulting corpus was tokenised into sentences and, to facilitate downstream data curation, we filtered for lines that explicitly mention the keywords, thus keeping the most descriptive lines for each gene. The LOF class was appended with genes annotated in the ClinGen database (Rehm et al. 2015) as “Sufficient evidence for dosage pathogenicity” (as of 2022-07-07), and a supporting evidence was added from the ClinGen entry. The raw evidence lines were manually reviewed and obvious false positives were removed. For example, in the LOF class a line may be: “… individuals harboring a heterozygous deletion in ATAD3A are unaffected suggesting a dominant-negative pathogenic mechanism or a gain-of-function mechanism for de novo missense variants rather than haploinsufficiency” (Harel et al. 2016), which explicitly dismisses haploinsufficiency as a molecular mechanism. Importantly, we noticed that a substantial fraction of GOF and DN evidence lines relate to artificial constructs used in biological research and had no association to human disease, which further necessitated manual curation. In overlap cases, when genes belong to multiple categories, we employed a hierarchical strategy to create a non-redundant gene list: DN > GOF > LOF. We made available the genes of different molecular mechanism classes in **Table S1**, containing also the evidence lines and the relevant PubMed identifiers.

### Protein functional classes

Functional classification of proteins were retrieved from PANTHER version 17.0 (Mi et al. 2021). In the category “Transporter” we included the classes “transporter”, “transmembrane signal receptor” and “membrane traffic protein”. In the category “Metabolic enzymes” we grouped “nucleic acid metabolism protein”, and “metabolite interconversion enzyme”. Finally, the category “TF/chromatin regulator” represents the combined classes of “gene-specific transcriptional regulator” and “chromatin/chromatin-binding, or -regulatory protein”.

### Protein abundance

Protein abundances were obtained from the integrated human dataset (version 2021) of PAXdb (Wang et al. 2015). Parts per million (ppm) values were converted to molar concentration based on the equation given by (Dubreuil, Matalon, and Levy 2019).

### HEK293 active ribosome profile

Normalised active ribosome protected fragments in the Human Embryonal Kidney 293 lineage were determined by (Clamer et al. 2018). The data is available via the NCBI Gene Expression Omnibus (Edgar, Domrachev, and Lash 2002) under accession GSE112353. Values were averaged over the two biological replicates and transcripts with values <1 were excluded from the analysis.

### Cotranslationally assembling proteins in HEK293 cells

The gene set was downloaded from the supplemental material of (Bertolini et al. 2021).

### Coiled coil motif containing proteins

Coiled coil motif containing proteins were retrieved from UniProt, using the search terms: (keyword:KW-0175) AND (organism_id:9606) AND (reviewed:true).

### Position of genes on chromosomes

Genes were mapped to chromosomes using the consensus coding sequence (CCDS) database (Pujar et al. 2018) downloaded via the NCBI FTP site.

### Molecular graphics

Visualisation of structures was performed with UCSF ChimeraX version 1.4 (Pettersen et al., 2021). The membrane embedded GABA_A_ receptor was sourced from the MemProtMD database (Newport, Sansom, and Stansfeld 2019).

### Statistical analyses

Data exploration and statistical analyses were carried out in RStudio (Rstudio 2022) “Spotted Wakerobin” release (2022-07-06), using R version 4.2.1 (R Development Core Team, 2021). The R packages used for analyses were: `tidyverse, tidytable, rsample, rstatix, scales, ggridges`, and ggbeeswarm. Error bars in bar charts represent 68% Jeffrey’s binomial confidence intervals and the probabilities between the proportions of cotranslationally assembling subunits were calculated from the hypergeometric distribution. The 95% confidence interval for the odds ratio was calculated with the standard error method (Bland and Altman 2000), where the value of the 97.5th percentile point of the normal distribution (∼1.96) was derived as `stats::qnorm(0.975)` in `R`. In Wilcoxon rank-sum tests the effect size was defined as the z-score computed from the p-value over the square root of sample size (Tomczak and Tomczak 2014). In multiple comparisons, the Holm–Bonferroni method (Holm, 1979) was used to correct for familywise error rate. In the bootstrap analysis, data were stratified for molecular mechanisms in 10,000 resamples. The p-value was calculated by determining the fraction of point estimates indicating a C-terminal interface shift, with correction for finite sampling (Davison and Hinkley 1997). The 95% confidence intervals of the bootstrap estimates were derived using the percentile method (Jung et al. 2019).

### Lasso regression – feature selection

To prioritise genes that mainly give rise to non-LOF mutations over those that harbour LOF mutations, the following variables were included in the model:

1. gnomAD mutational constraint metrics (Karczewski et al. 2020)
  - pLI – Probability that transcript falls into the distribution of haploinsufficient genes.
  - pRec – Probability that transcript falls into distribution of recessive genes.
  - oe_lof – Observed over expected ratio for predicted loss-of-function variants in transcript.
2. Sequence-derived or evolutionary variables
  - Protein-length – As per UniProt canonical isoform.
  - Number of paralogues (Huang et al. 2010) – Paralogues of human genes were called from Ensembl (Cunningham et al. 2022) via the biomaRt R package.
  - Maximum identity to paralogue (Huang et al. 2010) – We calculated the protein sequence identity of each gene to every one of its paralogues using Clustal Omega version 1.2.4 (Sievers et al. 2011) and the maximum identity was kept.
  - Human-macaque (*Macaca mulatta*) dN/dS (Huang et al. 2010) – The ratios of nonsynonymous to non-synonymous substitutions between human-macaque orthologues were called from Ensembl (Cunningham et al. 2022) via the biomaRt R package.
  - ncGERP++ – The phylogenetic conservation values of the genes’ regulatory sequences as derived from GERP++ scores were acquired from (Petrovski et al. 2015).
  - Number of domains (Huang et al. 2010) – The number of domains were taken from the human proteome-specific Pfam release 2021-11-15 (Mistry et al. 2021).
  - Membrane propensity – We calculated from protein sequences the mean value of the scale NAKH900110 “Normalized composition of membrane proteins” (Nakashima, Nishikawa, and Ooi 1990) from the AA index database (Kawashima et al. 2008).
3. Interaction network-based property
  - Betweenness centrality (Huang et al. 2010) – This measure was calculated from the human protein interaction network of the STRING database version 11.5 (Szklarczyk et al. 2021) at the default score threshold of 400 using the STRINGdb and igraph R packages.
4. Structural properties
  - Structural symmetry (monomer, heteromer, and homomeric symmetry groups: *C*_2_, *C*_n>2_, *D*_n>1_, and other homomeric symmetry), interface size and number of subunits.
  - Structure isoelectric point – as previously described in (Badonyi and Marsh 2022).
  - Absolute contact order (Plaxco, Simons, and Baker 1998) – We calculated the contact order from the AlphaFold predicted human structures using the perl script written by Eric Alm available at https://depts.washington.edu/bakerpg/contact_order/contactOrder.pl.
  - A copy of the script can be found in the OSF repository linked to this manuscript.
  - Mean structure pLDDT and alpha helix content.
5. Functional properties
  - Protein functional classification from this study. Proteins with functions other than those introduced earlier were classified as “known other function” and those lacking a functional annotation were classed as “unknown function”.
6. Experimental data
  - Protein abundance.
  - Cotranslational assembly annotations.
  - RNA expression variance – We accessed the rna_tissue_consensus.tsv.zip file from the Human Protein Atlas (Uhlen et al. 2010) on 2022-09-01 and calculated the variance in expression per gene across the 54 tissues.

### Lasso regression – data preparation

We assembled a dataset of 9,051 genes with the above features. Genes of monomers in the PDB were assigned an interface size of 0, a relative interface location of 0, a number of subunits of 1, and were assumed not to undergo cotranslational assembly even if they were detected by (Bertolini et al. 2021). Missing data in ten variables were imputed using five nearest neighbours (Gower 1971). The missing value rate were the following (per cent missing in brackets): ncGERP++ (9.7); human-macaque dN/dS (9.3); pLI, pRec, and oe_lof (7.3); betweenness centrality (5.6); protein abundance (4.2); number of domains (2.7); structure isoelectric point (2.1); RNA expression variance (1.3). Lastly, all nominal variables were one-hot encoded and numeric data was normalised to have a standard deviation of one and a mean of zero.

### Lasso regression – model building and performance evaluation

In an initial model screen, we evaluated the performance of more complex statistical learning methods, including a random forest, a support vector machine, and a single layer neural network. However, no performance gain was observed relative to a simpler and more interpretable regression model (data not shown). Logistic regression with lasso (least absolute shrinkage and selection operator) is a solution to fitting a model in which only certain variables play a role. The algorithm applies increasingly larger penalties to multivariable regression coefficients, shrinking those of less important variables to zero, causing their sequential drop-out (L1 regularisation) and thus retaining only informative features. First, to avoid inflating the model’s performance by presence of homologues, we performed redundancy filtering at 50% sequence identity on the full canonical protein sequences using the method described in the section **Relative interface location**. This procedure removed 88 from the 879 genes with experimentally available structure data and dominant molecular mechanism classifications. The remaining 791 genes, of which 543 (69%) are non-LOF and 248 (31%) are LOF, were split into 75% training and 25% test sets with 10-fold cross-validation performed on the training set and repeated 3 times. The model was tuned using 18 values of the λ parameter, generated to be log-linearly distributed between 0 and 1. The final value of λ = 0.00501 was chosen on the basis that it yielded the highest prediction accuracy in the assessment-folds of the cross-validation. The model was finalised on the entire training set and evaluated on the test set. The similarity of ROC AUCs measured on the cross-validation folds (0.735) versus on the test set (0.743) suggested that the model had not been overfitted. Variable importance was computed as the absolute values of the β coefficients scaled to the [0,1] interval. Model building and evaluation was performed using the `tidymodels R` metapackage. Thresholds T1 and T2 were derived using the `threshold_perf()` function from the R package probably.

## Data and code availability

Data and analysis code are available on OSF at https://osf.io/92uk5/.

## Acknowledgements

MB is supported by the Biotechnology and Biological Sciences Research Council EASTBIO Doctoral Training Programme (BB/M010996/1), and used resources provided by the Edinburgh Compute and Data Facility. JAM is supported by a European Research Council Consolidator Grant (101001169) and is a Lister Institute Research Prize Fellow.

---

**Figure S1.**
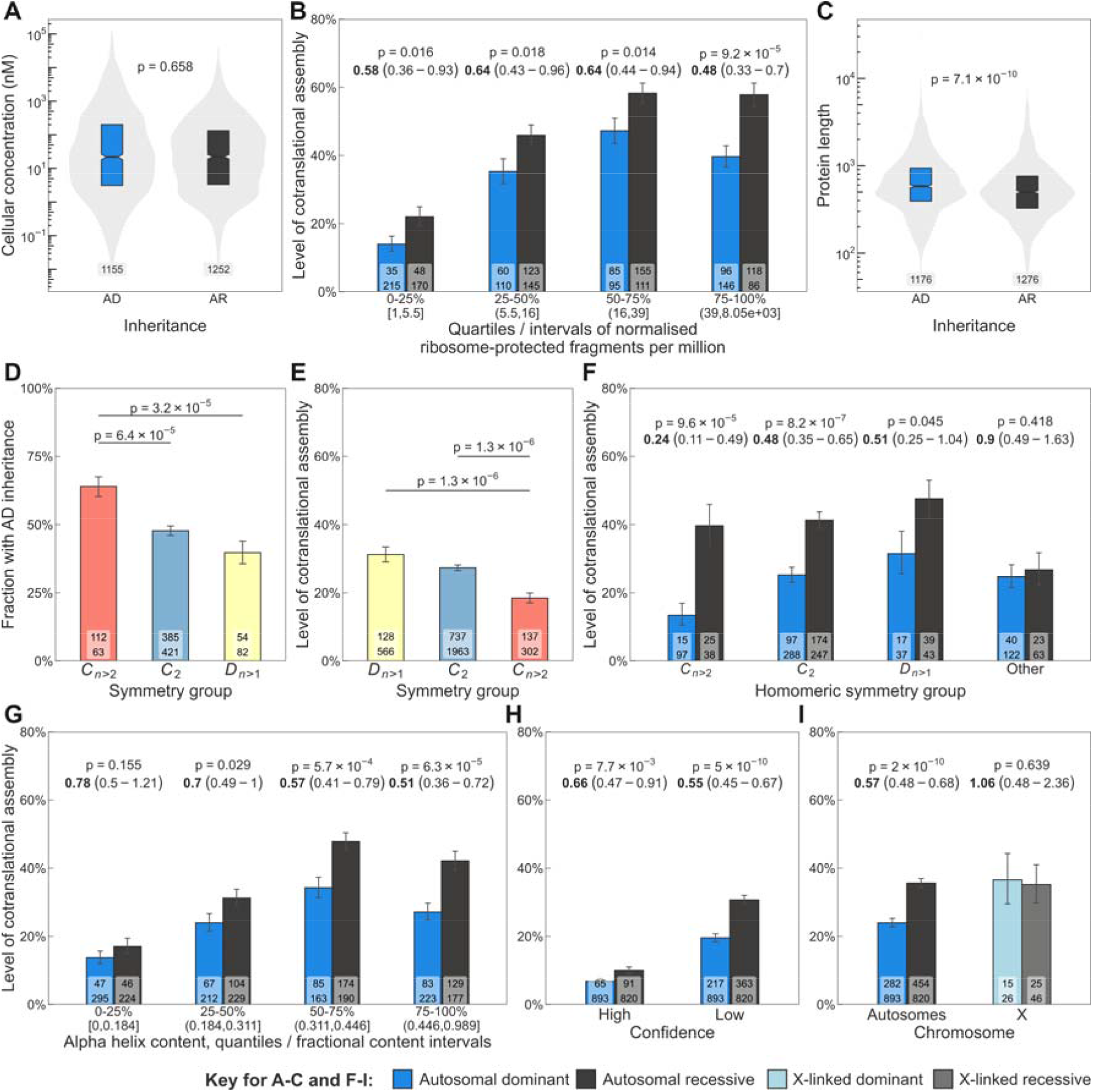
(**A**) Box-violin plot comparison of the abundance distribution of autosomal dominant (AD) and autosomal recessive (AR) homomers and repeated subunits. Boxes denote data within 25^th^ and 75^th^ percentiles, the middle line represents the median and the notch contains the 95% confidence interval of the median. Numbers at the bottom show sample size and the p-value was calculated with a Wilcoxon rank-sum test. (**B**) The level of cotranslational assembly split into quartiles of active ribosome protected fragment counts measured in HEK293 cells. Each bin corresponds to 25% of proteins by count and the fragment per million intervals are displayed in brackets. The p-value and odds ratio of the respective comparison is displayed above the bars. Count labels represent cotranslationally assembling (top) and all other proteins (bottom). (**C**) Box-violin plot comparison of the length distribution of AD and AR subunits. Numbers at the bottom represent sample size and the p-value was calculated with a Wilcoxon rank-sum test. (**D**) Fraction of homomeric symmetry groups with AD inheritance. Cyclic (*C*_n>2_), twofold (*C*_2_), dihedral (*D*_n>1_). The p-values are Holm-Bonferroni corrected and sampled from the hypergeometric distribution. Count labels represent AD (top) and AR (bottom) genes. (**E**) The level of cotranslational assembly among the main homomeric symmetry groups. The p-values are Holm-Bonferroni corrected and sampled from the hypergeometric distribution. Count labels represent cotranslationally assembling (top) and all other proteins (bottom). (**F**) Level of cotranslational assembly in genes of homomers with AD and AR disease inheritance binned by symmetry groups: cyclic (*C*n>2), twofold (*C*_2_), dihedral (*D*n>1) and other. (**G**) The level of cotranslational assembly split into quartiles of alpha helix content. Each bin corresponds to 25% of proteins by count and the fractional helix content intervals are displayed in brackets. (**H**) The level of cotranslational assembly split by the confidence in their identification. (**I**) Comparison of the level of cotranslational assembly in genes of subunits mapping to autosomes (AD vs AR) or to the X chromosome (X-linked dominant vs X-linked recessive).

**Figure S2.**
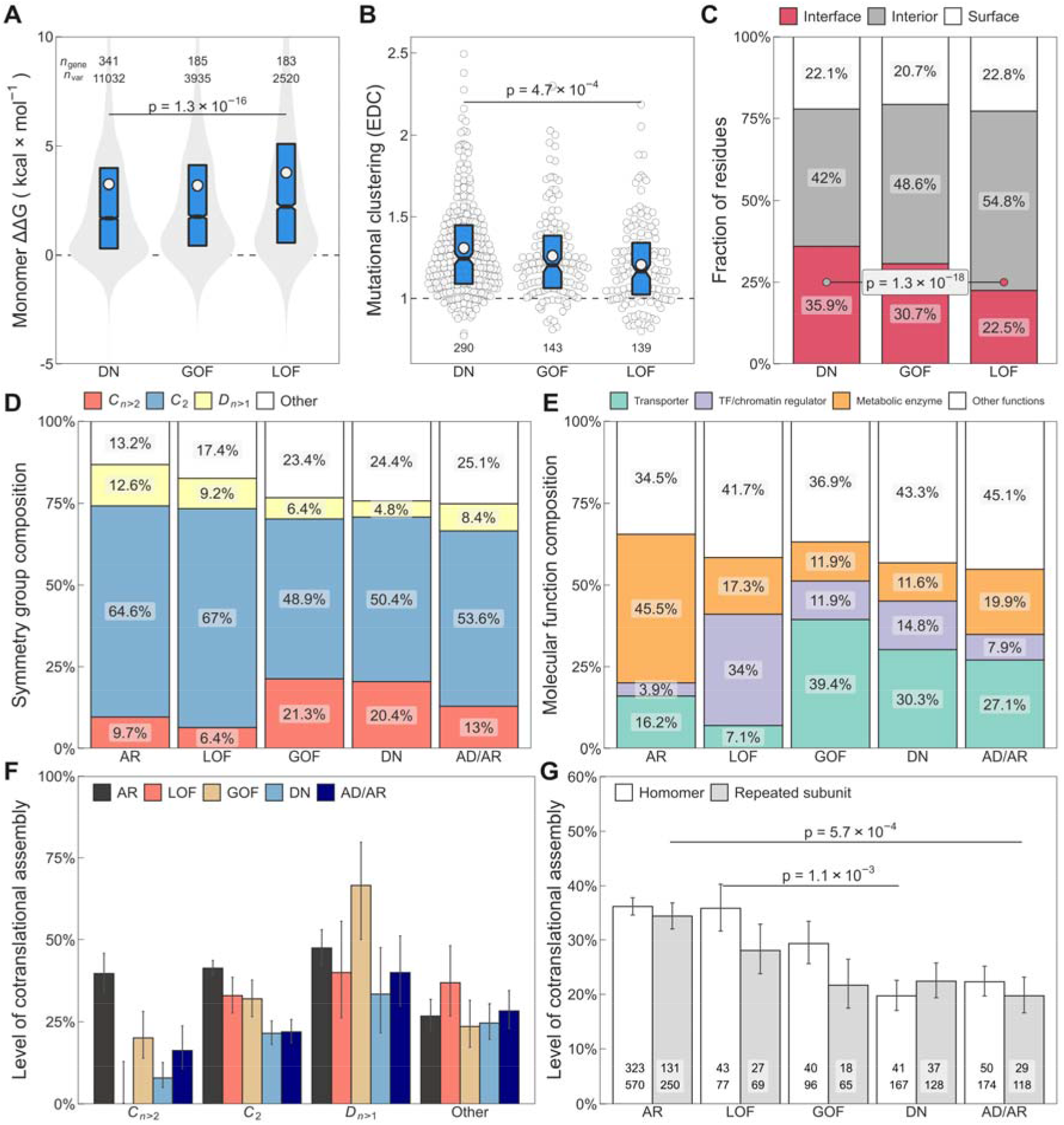
(**A**) Box-violin plot comparison of the predicted ΔΔG of pathogenic mutations in homomers and repeated subunits, grouped by molecular mechanism: dominant negative (DN), gain of function (GOF), and loss of function (LOF). Boxes denote data within 25^th^ and 75^th^ percentiles, the middle line represents the median, the notch contains the 95% confidence interval of the median and white dots are the mean. Numbers on top are sample sizes for genes (n_gene_) and missense variants (n_var_) within the groups. The p-value was calculated with a Wilcoxon rank-sum test. (**B**) Box-beeswarm plot comparison of the extent of disease clustering (EDC) metric that measures the extent to which pathogenic mutations cluster in 3D space. Numbers at the bottom denote the number of genes in each group. The p-value was calculated with a Wilcoxon rank-sum test. (**C**) Stacked bar chart showing the interface residue enrichment of missense pathogenic mutations in the DN group relative to LOF. The p-value was drawn from the hypergeometric distribution. (**D**) Stacked bar chart of the symmetry group composition of homomeric subunits with different inheritance and molecular mechanism: autosomal recessive (AR), mixed autosomal recessive / autosomal dominant (AD/AR), dominant negative (DN), gain of function (GOF), and loss of function (LOF). The symmetry groups are: cyclic (*C*_n>2_), twofold (*C*_2_), dihedral (*D*_n>1_), and other. (**E**) Stacked bar chart of the molecular function composition of homomers and repeated subunit heteromers with different inheritance and molecular mechanism. (**F**) Level of cotranslational assembly within the different inheritance and molecular mechanism classes subset by homomeric symmetry groups. (**H**) Level of cotranslational assembly split into homomers and repeated subunit heteromers. The p-values were calculated from the hypergeometric distribution.

**Figure S3.**
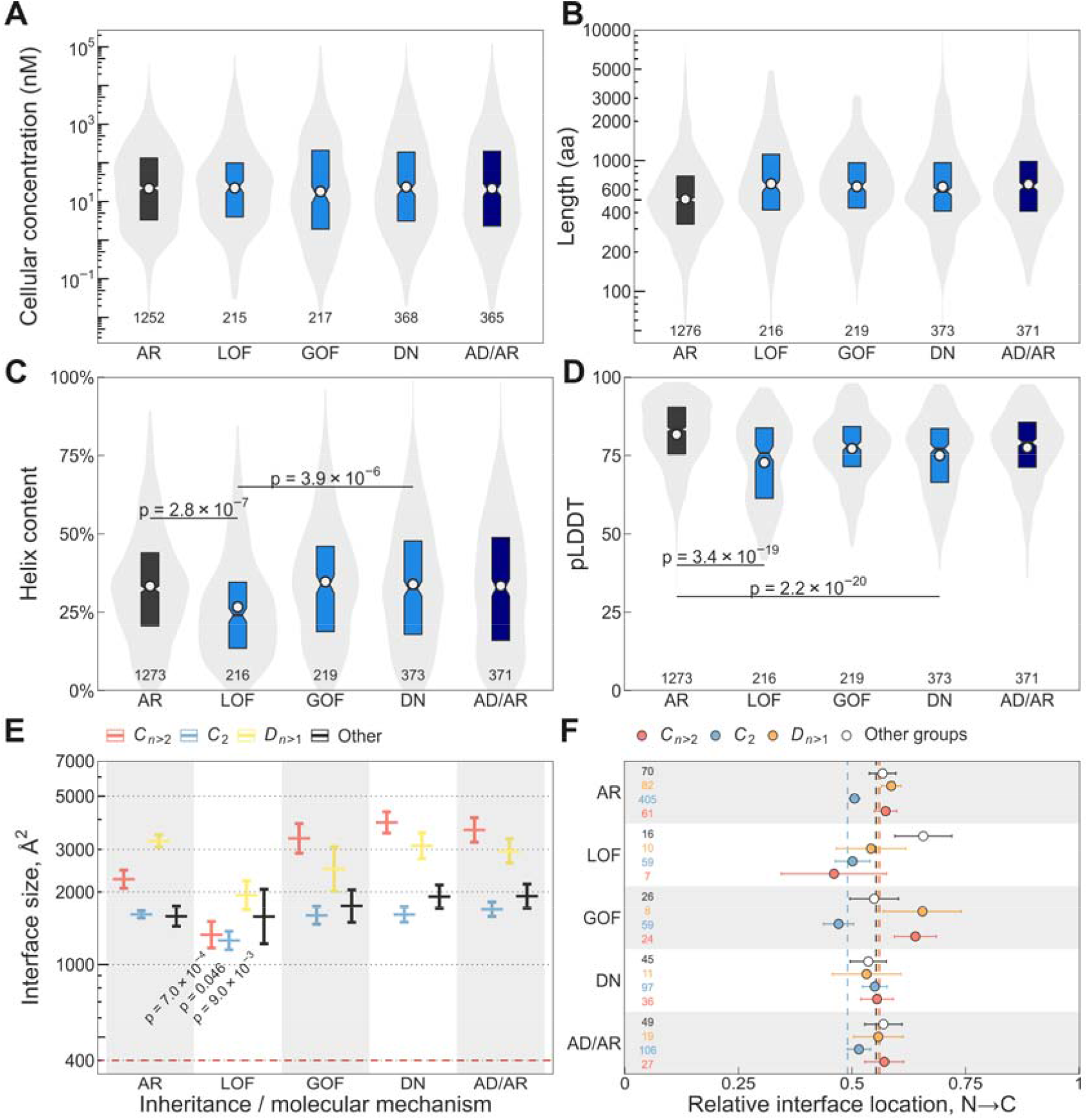
(**A**) Box-violin plot comparison of the abundance, length (**B**), helix content (**C**), and pLDDT (**D**) distribution of homomers and repeated subunits with different inheritance and molecular mechanism: autosomal recessive (AR), mixed autosomal recessive / autosomal dominant (AD/AR), dominant negative (DN), gain of function (GOF), and loss of function (LOF). Boxes denote data within 25^th^ and 75^th^ percentiles, the middle line represents the median, the notch contains the 95% confidence interval of the median and white dots are the mean. The p-values are from a Holm-Bonferroni corrected Dunn’s test of multiple comparison. Sample sizes are indicated at the bottom of each violin. (**E**) Interface area differences in homomers with different inheritance and molecular mechanism. The homomeric symmetry groups are: cyclic (*C*n>2), twofold (*C*_2_), dihedral (*D*n>1), and other. Crossbars are mean ± SEM. The p-values indicate differences between LOF vs DN and are from Holm-Bonferroni corrected Dunn’s test of multiple comparison. Sample sizes are shown on panel (**F**). (**F**) Relative interface location of homomers with different inheritance and molecular mechanism. Dashed lines correspond to the population mean of the corresponding symmetry group, indicated by its colour. Sample sizes are shown on the left.

**Figure S4.**
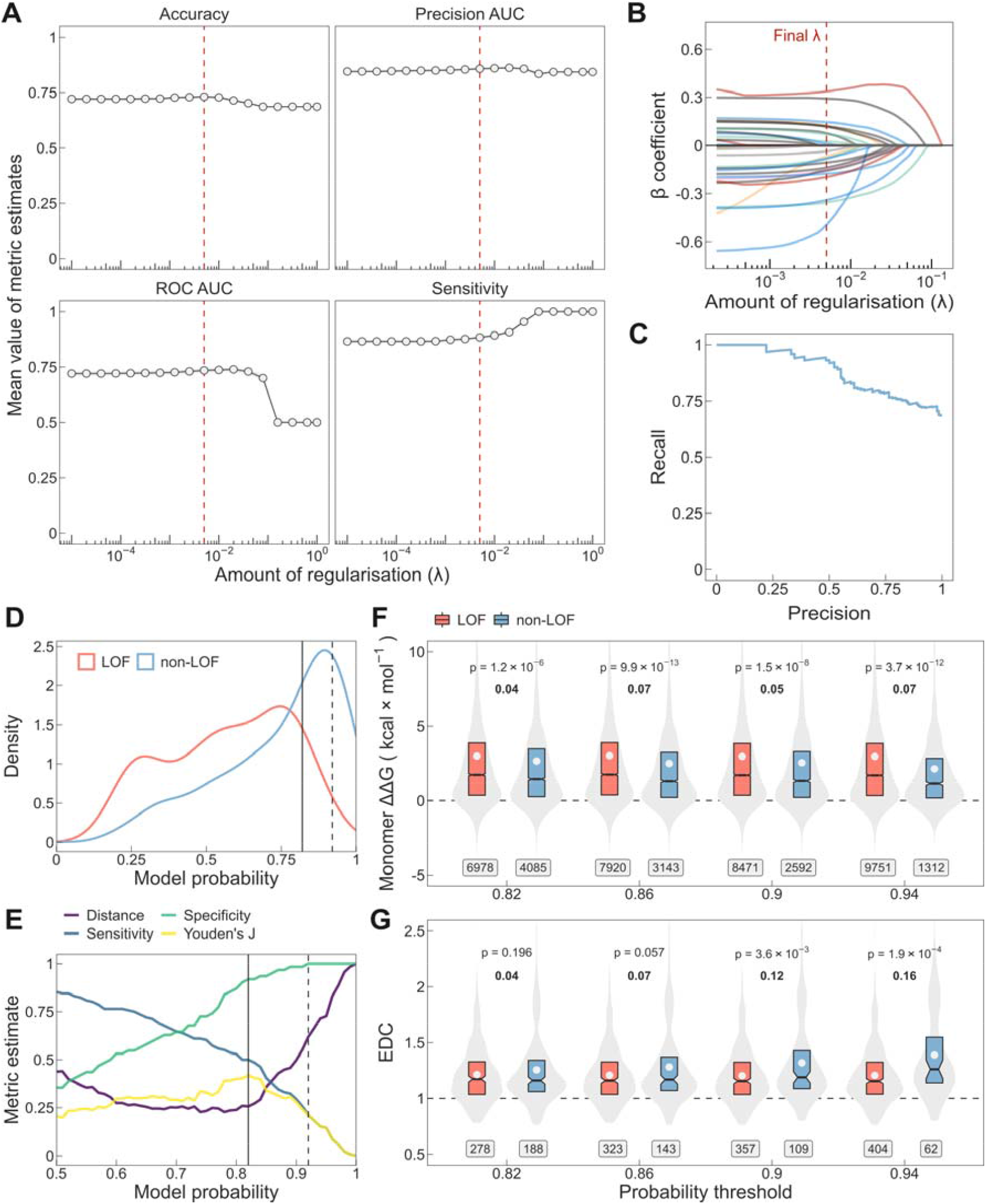
(**A**) Performance of the lasso regression model as a function of the penalty parameter estimated from the cross-validation folds. Dashed line is at λ = 0.00501, the penalty value in the final model. (**B**) Lasso penalty (λ) vs the regression coefficient (β). The lines are coloured according to the type of the variable: sequence-derived or evolutionary variables (blue), functional annotations (green), mutational constraint metrics (red), structural properties (black), interaction network-based property (pink), and experimental data (orange). (**C**) Precision-recall curve of the lasso regression model measured on the test set. (**D**) Distribution of model probabilities for non-LOF and LOF genes in the test set. The solid vertical line marks the recommended threshold p = 0.82 (T1), the probability at which Youden’s J statistic has its maximum value. The dashed line marks p = 0.92 (T2), the probability at which the model reaches maximum specificity. (**E**) Performance metrics measured on the test set as a function of the model probability threshold. The distance is defined as (1 - sensitivity) ^ 2 + (1 - specificity) ^ 2 and Youden’s J statistic as sensitivity + specificity - 1. Solid and dashed vertical lines are defined in (**D**). (**F**) Differences in Gibbs free energy change and the extent of disease clustering (EDC) (**G**) of pathogenic mutations between proteins predicted to be non-LOF versus all other proteins measured at different thresholds. Genes that were used for training the model as well as known autosomal recessive genes were excluded. Boxes denote data within 25^th^ and 75^th^ percentiles with the middle line representing the median, the notch contains the 95% confidence interval of the median and the white dots are the mean. Labels indicate the number of variants (ΔΔG) or the number of proteins (EDC) in the groups. The p-values were calculated with Wilcoxon rank-sum tests and effect sizes are shown in bold.

**Table S1** Autosomal dominant genes and their molecular mechanism classifications, with supporting evidence lines and their respective PubMed identifier.

**Table S2** Predicted non-LOF probabilities for human genes from the lasso regression model. Additional columns indicate if the gene was part of the training set, or if the gene has known dominant or recessive associations recorded in OMIM.

## References

Akdel, Mehmet et al. 2022. “A Structural Biology Community Assessment of AlphaFold2 Applications.” Nature Structural & Molecular Biology: 1–12.

Amberger, Joanna S. et al. 2015. “OMIM.Org: Online Mendelian Inheritance in Man (OMIM®), an Online Catalog of Human Genes and Genetic Disorders.” Nucleic Acids Research 43(D1): D789–98.

Attard, Thomas J., Julie P. I. Welburn, and Joseph A. Marsh. 2022. “Understanding Molecular Mechanisms and Predicting Phenotypic Effects of Pathogenic Tubulin Mutations.” PLOS Computational Biology 18(10): e1010611.

Backwell, Lisa, and Joseph A. Marsh. 2022. “Diverse Molecular Mechanisms Underlying Pathogenic Protein Mutations: Beyond the Loss-of-Function Paradigm.” Annual review of genomics and human genetics 23(1).

Badonyi, Mihaly, and Joseph A Marsh. 2022. “Large Protein Complex Interfaces Have Evolved to Promote Cotranslational Assembly.” eLife 11. https://elifesciences.org/articles/79602.

Bergendahl, L. Therese et al. 2019. “The Role of Protein Complexes in Human Genetic Disease.” Protein Science 28(8): 1400–1411.

Bergendahl, L. Therese, and Joseph A. Marsh. 2017. “Functional Determinants of Protein Assembly into Homomeric Complexes.” Scientific Reports 7(1).

Berman, Helen M. et al. 2000. “The Protein Data Bank.” Nucleic Acids Research.

Bertolini, Matilde et al. 2021. “Interactions between Nascent Proteins Translated by Adjacent Ribosomes Drive Homomer Assembly.” Science 371(6524): 57–64.

Birgmeier, Johannes et al. 2020. “AMELIE Speeds Mendelian Diagnosis by Matching Patient Phenotype and Genotype to Primary Literature.” Science Translational Medicine 12(544): eaau9113.

Bland, J. Martin, and Douglas G. Altman. 2000. “The Odds Ratio.” BMJ 320(7247): 1468.

Bowne, Sara J. et al. 2011. “A Dominant Mutation in RPE65 Identified by Whole-Exome Sequencing Causes Retinitis Pigmentosa with Choroidal Involvement.” European Journal of Human Genetics 19(10): 1074–81.

Cavaco, Branca M et al. 2018. “Homozygous Calcium-Sensing Receptor Polymorphism R544Q Presents as Hypocalcemic Hypoparathyroidism.” The Journal of Clinical Endocrinology & Metabolism 103(8): 2879–88.

Celesia, Gastone G. 2001. “Disorders of Membrane Channels or Channelopathies.” Clinical Neurophysiology 112(1): 2–18.

Chen, Xiuzhen, and Christine Mayr. 2022. “A Working Model for Condensate RNA-Binding Proteins as Matchmakers for Protein Complex Assembly.” RNA 28(1): 76–87.

Chicco, Davide, Niklas Tötsch, and Giuseppe Jurman. 2021. “The Matthews Correlation Coefficient (MCC) Is More Reliable than Balanced Accuracy, Bookmaker Informedness, and Markedness in Two-Class Confusion Matrix Evaluation.” BioData Mining 14(1): 13.

Clamer, Massimiliano et al. 2018. “Active Ribosome Profiling with RiboLace.” Cell Reports 25(4): 1097-1108.e5.

Cunningham, Fiona et al. 2022. “Ensembl 2022.”Nucleic Acids Research 50(D1): D988–95.

Curtis, Andrew R.J. et al. 2001. “Mutation in the Gene Encoding Ferritin Light Polypeptide Causes Dominant Adult-Onset Basal Ganglia Disease.” Nature Genetics 28(4): 350–54.

Dang, Vinh T., Karin S. Kassahn, Andrés Esteban Marcos, and Mark A. Ragan. 2008. “Identification of Human Haploinsufficient Genes and Their Genomic Proximity to Segmental Duplications.” European Journal of Human Genetics 2008 16:11 16(11): 1350–57.

Davison, A. C., and D. V. Hinkley. 1997. Bootstrap Methods and Their Application. Cambridge: Cambridge University Press.

Delgado, Javier, Leandro G. Radusky, Damiano Cianferoni, and Luis Serrano. 2019. “FoldX 5.0: Working with RNA, Small Molecules and a New Graphical Interface.” Bioinformatics 35(20): 4168–69.

Demircioglu, F. Esra et al. 2019. “The AAA□+□ATPase TorsinA Polymerizes into Hollow Helical Tubes with 8.5 Subunits per Turn.” Nature Communications 10(1): 3262.

Dénes, Türei et al. 2021. “Integrated Intra- and Intercellular Signaling Knowledge for Multicellular Omics Analysis.” Molecular Systems Biology 17(3): e9923.

Drutman, Scott B. et al. 2019. “Homozygous NLRP1 Gain-of-Function Mutation in Siblings with a Syndromic Form of Recurrent Respiratory Papillomatosis.” Proceedings of the National Academy of Sciences 116(38): 19055–63.

Dubreuil, Benjamin, Or Matalon, and Emmanuel D. Levy. 2019. “Protein Abundance Biases the Amino Acid Composition of Disordered Regions to Minimize Non-Functional Interactions.” Journal of Molecular Biology 431(24): 4978–92.

Edgar, Ron, Michael Domrachev, and Alex E. Lash. 2002. “Gene Expression Omnibus: NCBI Gene Expression and Hybridization Array Data Repository.” Nucleic acids research 30(1): 207–10.

Eilbeck, Karen, Aaron Quinlan, and Mark Yandell. 2017. “Settling the Score: Variant Prioritization and Mendelian Disease.” Nature Reviews Genetics 2017 18:10 18(10): 599–612.

Forrest, Lucy R. 2015. “Structural Symmetry in Membrane Proteins∗.” Annual Review of Biophysics 44(1): 311–37.

Geller, David S. et al. 2000. “Activating Mineralocorticoid Receptor Mutation in Hypertension Exacerbated by Pregnancy.” Science 289(5476): 119–23.

Gerasimavicius, Lukas, Benjamin J. Livesey, and Joseph A. Marsh. 2022. “Loss-of-Function, Gain-of-Function and Dominant-Negative Mutations Have Profoundly Different Effects on Protein Structure.” Nature Communications 2022 13:1 13(1): 1–15.

Gerber, Sylvie et al. 2016. “Recessive and Dominant De Novo ITPR1 Mutations Cause Gillespie Syndrome.” American Journal of Human Genetics 98(5): 971–80.

Gilmore, Ross et al. 1996. “Co-Translational Trimerization of the Reovirus Cell Attachment Protein.” EMBO Journal 15(11): 2651–58.

Goodsell, David S., and Arthur J. Olson. 2000. “Structural Symmetry and Protein Function.” Annual Review of Biophysics and Biomolecular Structure 29(1): 105–53.

Gower, J. C. 1971. “A General Coefficient of Similarity and Some of Its Properties.” Biometrics 27(4): 857– 71.

Haldane, J. B. S. 1930. “A Note on Fisher’s Theory of the Origin of Dominance, and on a Correlation between Dominance and Linkage.” The American Naturalist 64(690): 87–90.

Harel, Tamar et al. 2016. “Recurrent De Novo and Biallelic Variation of ATAD3A, Encoding a Mitochondrial Membrane Protein, Results in Distinct Neurological Syndromes.” American Journal of Human Genetics 99(4): 831–45.

Hochberg, Georg K.A. et al. 2020. “A Hydrophobic Ratchet Entrenches Molecular Complexes.” Nature 588(7838): 503–8.

Holm, Sture. 1979. “A Simple Sequentially Rejective Multiple Test Procedure.” Scandinavian Journal of Statistics 6(2).

Horvath, Gabriella A. et al. 2018. “Gain-of-Function KCNJ6 Mutation in a Severe Hyperkinetic Movement Disorder Phenotype.” Neuroscience 384: 152–64.

Huang, Ni, Insuk Lee, Edward M. Marcotte, and Matthew E. Hurles. 2010. “Characterising and Predicting Haploinsufficiency in the Human Genome” ed. Mikkel H. Schierup. PLoS Genetics 6(10): e1001154.

Hurst, Laurence D., and James P. Randerson. 2000. “Dosage, Deletions and Dominance: Simple Models of the Evolution of Gene Expression.” Journal of Theoretical Biology 205(4): 641–47.

Ishii, Atsushi et al. 2017. “A de Novo Missense Mutation of GABRB2 Causes Early Myoclonic Encephalopathy.” Journal of Medical Genetics 54(3): 202–11.

Jimenez-Sanchez, G., B. Childs, and D. Valle. 2001. “Human Disease Genes.” Nature 409(6822): 853–55.

Jin, Weili, Hidnori Takagi, Bruno Pancorbo, and Elizabeth C Theil. 2001. “‘Opening’ the Ferritin Pore for Iron Release by Mutation of Conserved Amino Acids at Interhelix and Loop Sites †.”

Johnston, Iain G. et al. 2022. “Symmetry and Simplicity Spontaneously Emerge from the Algorithmic Nature of Evolution.” Proceedings of the National Academy of Sciences 119(11): e2113883119.

Jung, Kwanghee, Jaehoon Lee, Vibhuti Gupta, and Gyeongcheol Cho. 2019. “Comparison of Bootstrap Confidence Interval Methods for GSCA Using a Monte Carlo Simulation.” Frontiers in Psychology 10. https://www.frontiersin.org/articles/10.3389/fpsyg.2019.02215 (August 30, 2022).

Kabsch, Wolfgang, and Christian Sander. 1983. “Dictionary of Protein Secondary Structure: Pattern Recognition of Hydrogen-bonded and Geometrical Features.” Biopolymers 22(12): 2577–2637.

Kacser, H., and J. A. Burns. 1981. “The Molecular Basis of Dominance.” Genetics 97(3–4): 639–66.

Kamenova, Ivanka et al. 2019. “Co-Translational Assembly of Mammalian Nuclear Multisubunit Complexes.” Nature Communications 10(1).

Karczewski, Konrad J. et al. 2020. “The Mutational Constraint Spectrum Quantified from Variation in 141,456 Humans.” Nature 581(7809): 434–43.

Kawashima, Shuichi et al. 2008. “AAindex: Amino Acid Index Database, Progress Report 2008.” Nucleic Acids Research 36(suppl_1): D202–5.

Kiser, Philip D. et al. 2015. “Catalytic Mechanism of a Retinoid Isomerase Essential for Vertebrate Vision.” Nature chemical biology 11(6): 409–15.

Lelieveld, Stefan H. et al. 2017. “Spatial Clustering of de Novo Missense Mutations Identifies Candidate Neurodevelopmental Disorder-Associated Genes.” American Journal of Human Genetics 101(3): 478–84.

Liebeskind, Benjamin J., David M. Hillis, and Harold H. Zakon. 2015. “Convergence of Ion Channel Genome Content in Early Animal Evolution.” Proceedings of the National Academy of Sciences of the United States of America 112(8): E846–51.

Liu, Jiangang et al. 2006. “Intrinsic Disorder in Transcription Factors.” Biochemistry 45(22): 6873–88.

Lomize, Andrei L. et al. 2022. “Membranome 3.0: Database of Single-Pass Membrane Proteins with AlphaFold Models.” Protein Science 31(5): e4318.

Lynch, Michael. 2013. “Evolutionary Diversification of the Multimeric States of Proteins.” Proceedings of the National Academy of Sciences of the United States of America 110(30): E2821–28.

Ma, Weirui, and Christine Mayr. 2018. “A Membraneless Organelle Associated with the Endoplasmic Reticulum Enables 3^′^UTR-Mediated Protein-Protein Interactions.” Cell 175(6): 1492-1506.e19.

MacArthur, Daniel G. et al. 2012. “A Systematic Survey of Loss-of-Function Variants in Human Protein-Coding Genes.” Science 335(6070): 823–28.

Mallik, Saurav, Dan S Tawfik, and Emmanuel D Levy. 2022. “How Gene Duplication Diversifies the Landscape of Protein Oligomeric State and Function.” Current Opinion in Genetics & Development 76: 101966.

Marsh, Joseph A., Holly A. Rees, Sebastian E. Ahnert, and Sarah A. Teichmann. 2015. “Structural and Evolutionary Versatility in Protein Complexes with Uneven Stoichiometry.” Nature Communications 6(1): 1–10.

Marsh, Joseph A, and Sarah A Teichmann. 2015. “Structure, Dynamics, Assembly, and Evolution of Protein Complexes.” Annual Review of Biochemistry 84: 551–75.

McEntagart, Meriel et al. 2016. “A Restricted Repertoire of de Novo Mutations in ITPR1 Cause Gillespie Syndrome with Evidence for Dominant-Negative Effect.” American Journal of Human Genetics 98(5): 981–92.

Mi, Huaiyu et al. 2021. “PANTHER Version 16: A Revised Family Classification, Tree-Based Classification Tool, Enhancer Regions and Extensive API.” Nucleic Acids Research 49(D1): D394–403.

Mistry, Jaina et al. 2021. “Pfam: The Protein Families Database in 2021.” Nucleic Acids Research 49(D1): D412–19.

Morales-Polanco, Fabian et al. 2021. “Core Fermentation (CoFe) Granules Focus Coordinated Glycolytic MRNA Localization and Translation to Fuel Glucose Fermentation.” iScience 24(2): 102069.

Morales-Polanco, Fabián, Jae Ho Lee, Natália M. Barbosa, and Judith Frydman. 2022. “Cotranslational Mechanisms of Protein Biogenesis and Complex Assembly in Eukaryotes.” Annual Review of Biomedical Data Science 5: 67–94.

Mrazek, Jan et al. 2014. “Polyribosomes Are Molecular 3D Nanoprinters That Orchestrate the Assembly of Vault Particles.” ACS Nano 8(11): 11552–59.

Nakashima, Hiroshi, Ken Nishikawa, and Tatsuo Ooi. 1990. “Distinct Character in Hydrophobicity of Amino Acid Compositions of Mitochondrial Proteins.” Proteins: Structure, Function, and Bioinformatics 8(2): 173–78.

Natan, Eviatar et al. 2018. “Cotranslational Protein Assembly Imposes Evolutionary Constraints on Homomeric Proteins.” Nature Structural and Molecular Biology 25(3): 279–88.

Natan, Eviatar, Jonathan N. Wells, Sarah A. Teichmann, and Joseph A. Marsh. 2017. “Regulation, Evolution and Consequences of Cotranslational Protein Complex Assembly.” Current Opinion in Structural Biology 42: 90–97.

Newport, Thomas D, Mark S P Sansom, and Phillip J Stansfeld. 2019. “The MemProtMD Database: A Resource for Membrane-Embedded Protein Structures and Their Lipid Interactions.” Nucleic Acids Research 47(D1): D390–97.

Nicholls, Chris D, Kevin G Mclure, Michael A Shields, and Patrick W K Lee. 2002. “Biogenesis of P53 Involves Cotranslational Dimerization of Monomers and Posttranslational Dimerization of Dimers.” Journal of Biological Chemistry 277(15): 12937–45.

Paila, Umadevi, Brad A. Chapman, Rory Kirchner, and Aaron R. Quinlan. 2013. “GEMINI: Integrative Exploration of Genetic Variation and Genome Annotations.” PLOS Computational Biology 9(7): e1003153.

Panasenko, Olesya O. et al. 2019. “Co-Translational Assembly of Proteasome Subunits in NOT1-Containing Assemblysomes.” Nature Structural and Molecular Biology 26(2): 110–20.

Peeters, Carel F. W. et al. 2019. “Stable Prediction with Radiomics Data.” http://arxiv.org/abs/1903.11696.

Perica, Tina et al. 2012. “The Emergence of Protein Complexes: Quaternary Structure, Dynamics and Allostery.” In Biochemical Society Transactions, Biochem Soc Trans, 475–91.

Petrovski, Slavé et al. 2015. “The Intolerance of Regulatory Sequence to Genetic Variation Predicts Gene Dosage Sensitivity” ed. Chris Cotsapas. PLOS Genetics 11(9): e1005492.

Plaxco, K. W., K. T. Simons, and D. Baker. 1998. “Contact Order, Transition State Placement and the Refolding Rates of Single Domain Proteins.” Journal of Molecular Biology 277(4): 985–94.

Prasad Bahadur Ranjit, Pinak Chakrabarti, Francis Rodier, and Joël Janin. 2004. “A Dissection of Specific and Non-Specific Protein–Protein Interfaces.” Journal of Molecular Biology 336(4): 943–55.

Pujar, Shashikant et al. 2018. “Consensus Coding Sequence (CCDS) Database: A Standardized Set of Human and Mouse Protein-Coding Regions Supported by Expert Curation.” Nucleic Acids Research 46(D1): D221–28.

Redick, S.D., and J.E. Schwarzbauer. 1995. “Rapid Intracellular Assembly of Tenascin Hexabrachions Suggests a Novel Cotranslational Process.” Journal of Cell Science 108(4): 1761–69.

Rehm, Heidi L. et al. 2015. “ClinGen — The Clinical Genome Resource.” New England Journal of Medicine 372(23): 2235–42.

Rstudio, Team. 2022. Rstudio Team, PBC, Boston, MA URL http://www.rstudio.com/ RStudio: Integrated Development for R.

Sayers, Eric W et al. 2021. “Database Resources of the National Center for Biotechnology Information.” Nucleic Acids Research.

Schwarz, Andre, and Martin Beck. 2019. “The Benefits of Cotranslational Assembly: A Structural Perspective.” Trends in Cell Biology 29(10): 791–803.

Seidel, Maximilian et al. 2022. “Co-Translational Assembly Orchestrates Competing Biogenesis Pathways.” Nature Communications 13(1): 1–15.

Seidman, J. G., and Christine Seidman. 2002. “Transcription Factor Haploinsufficiency: When Half a Loaf Is Not Enough.” The Journal of Clinical Investigation 109(4): 451–55.

Shiber, Ayala et al. 2018. “Cotranslational Assembly of Protein Complexes in Eukaryotes Revealed by Ribosome Profiling.” Nature 561(7722): 268–72.

Shihab, Hashem A., Mark F. Rogers, Colin Campbell, and Tom R. Gaunt. 2017. “HIPred: An Integrative Approach to Predicting Haploinsufficient Genes.” Bioinformatics (Oxford, England) 33(12): 1751– 57.

Sievers, Fabian et al. 2011. “Fast, Scalable Generation of High-Quality Protein Multiple Sequence Alignments Using Clustal Omega.” Molecular Systems Biology 7(1): 539.

Steinberg, Julia, Frantisek Honti, Stephen Meader, and Caleb Webber. 2015. “Haploinsufficiency Predictions without Study Bias.” Nucleic Acids Research 43(15): e101.

Szklarczyk, Damian et al. 2021. “The STRING Database in 2021: Customizable Protein–Protein Networks, and Functional Characterization of User-Uploaded Gene/Measurement Sets.” Nucleic Acids Research 49(D1): D605–12.

Thompson, Debra A. et al. 2002. “Retinal Dystrophy Due to Paternal Isodisomy for Chromosome 1 or Chromosome 2, with Homoallelism for Mutations in RPE65 or MERTK, Respectively.” American Journal of Human Genetics 70(1): 224–29.

Tomczak, Maciej, and Ewa Tomczak. 2014. “The Need to Report Effect Size Estimates Revisited. An Overview of Some Recommended Measures of Effect Size.” TRENDS in Sport Sciences 1(21): 19–25.

Torres, Gonzalo E. et al. 2004. “Effect of TorsinA on Membrane Proteins Reveals a Loss of Function and a Dominant-Negative Phenotype of the Dystonia-Associated DeltaE-TorsinA Mutant.” Proceedings of the National Academy of Sciences of the United States of America 101(44): 15650–55.

Tukiainen, Taru et al. 2017. “Landscape of X Chromosome Inactivation across Human Tissues.” Nature 550(7675): 244–48.

Tunyasuvunakool, Kathryn et al. 2021. “Highly Accurate Protein Structure Prediction for the Human Proteome.” Nature 596(7873): 590–96.

Uhlen, Mathias et al. 2010. “Towards a Knowledge-Based Human Protein Atlas.” Nature Biotechnology 28(12): 1248–50.

Wang, Gao T., Bo Peng, and Suzanne M. Leal. 2014. “Variant Association Tools for Quality Control and Analysis of Large-Scale Sequence and Genotyping Array Data.” The American Journal of Human Genetics 94(5): 770–83.

Wang, Kai, Mingyao Li, and Hakon Hakonarson. 2010. “ANNOVAR: Functional Annotation of Genetic Variants from High-Throughput Sequencing Data.” Nucleic Acids Research 38(16): e164.

Wang, Mingcong et al. 2015. “Version 4.0 of PaxDb: Protein Abundance Data, Integrated across Model Organisms, Tissues, and Cell-Lines.” Proteomics 15(18): 3163–68.

Wishner, B. C., K. B. Ward, E. E. Lattman, and W. E. Love. 1975. “Crystal Structure of Sickle-Cell Deoxyhemoglobin at 5 Å Resolution.” Journal of Molecular Biology 98(1): 179–94.

Wright, Sewall. 1929. “Fisher’s Theory of Dominance.” The American Naturalist 63(686): 274–79.

Wright, Sewall. 1934. “Physiological and Evolutionary Theories of Dominance.” The American Naturalist 68(714): 24–53.

Youden, W. J. 1950. “Index for Rating Diagnostic Tests.” Cancer 3(1): 32–35.

Zemojtel, Tomasz et al. 2014. “Effective Diagnosis of Genetic Disease by Computational Phenotype Analysis of the Disease-Associated Genome.” Science Translational Medicine 6(252): 252ra123–252ra123.

